# Monopogen: single nucleotide variant calling from single cell sequencing

**DOI:** 10.1101/2022.12.04.519058

**Authors:** Jinzhuang Dou, Yukun Tan, Kian Hong Kock, Jun Wang, Xuesen Cheng, Le Min Tan, Kyung Yeon Han, Chung Chau Hon, Woong Yang Park, Jay W Shin, Han Chen, Shyam Prabhakar, Nicholas Navin, Rui Chen, Ken Chen

## Abstract

Distinguishing how genetics impact cellular processes can improve our understanding of variable risk for diseases. Although single-cell omics have provided molecular characterization of cell types and states on diverse tissue samples, their genetic ancestry and effects on cellular molecular traits are largely understudied. Here, we developed Monopogen, a computational tool enabling researchers to detect single nucleotide variants (SNVs) from a variety of single cell transcriptomic and epigenomic sequencing data. It leverages linkage disequilibrium from external reference panels to identify germline SNVs from sparse sequencing data and uses Monovar to identify novel SNVs at cluster (or cell type) levels. Monopogen can identify 100K~3M germline SNVs from various single cell sequencing platforms (scRNA-seq, snRNA-seq, snATAC-seq etc), with genotyping accuracy higher than 95%, when compared against matched whole genome sequencing data. We applied Monopogen on human retina, normal breast and Asian immune diversity atlases, showing that that derived genotypes enable accurate global and local ancestry inference and identification of admixed samples from ancestrally diverse donors. In addition, we applied Monopogen on ~4M cells from 65 human heart left ventricle single cell samples and identified novel variants associated with cardiomyocyte metabolic levels and epigenomic programs. In summary, Monopogen provides a novel computational framework that brings together population genetics and single cell omics to uncover genetic determinants of cellular quantitative traits.

## Introduction

Defining the precise cellular contexts in which risk-associated variants affect cellular processes will help to better understand the molecular mechanisms of disease risks and to inform therapeutic strategies. This is important because recent studies have shown that many genetic variants affect tissue traits in a cell-type specific manner [1, 2]. Traditional bulk RNA analysis are usually biased towards abundant cell types defined by a limited set of marker genes [3].

Single cell sequencing has enabled comprehensive estimation of cellular composition and acquisition of cell type-specific molecular profiles [4], including rare cell types [5]. As opposed to bulk data, single-cell data allow linking genetics to cellular molecular traits such as variability in cellular expressions [6], cell type abundance [7], and gene regulatory networks [8]. As such, single-cell analyses in a population-based setting are becoming mainstream [9].

Although single-cell omics projects are increasingly profiling cell types/states on diverse tissue samples, such as those collected by the ancestry networks of human cell atlas (HCA) [10] and human tumor atlas network (HTAN) [11], the genetic ancestry of the samples and its contribution to cellular molecular traits are largely unexplored. To acquire an accurate genetic profile, it is often necessary to re-sequence the study samples using bulk whole genome/exome sequencing, which requires additional sequencing efforts and costs.

A potential cost-effective approach is to call genetic variants directly from single cell sequencing data, akin to previous studies using low-pass whole genome sequencing (WGS) [12] or bulk RNA sequencing [13]. A systematic comparison shows that traditional tools for bulk analysis, such as Samools [14], GATK [15], etc., detected less than 8% of variants from full-length SMART-seq2 data and considerably less from droplet-based data [16]. Possible reasons for low variant detection include: 1) the single cell RNA sequencing reads are usually enriched in specific genomic regions, such as 5’ or 3’ end of genes; 2) genes are usually expressed in cell type/state-specific patterns, and thus are highly variable across genome regions, leading to uneven sequencing depth distribution; 3) coverage is likely affected by allelic imbalance inherent in RNA profiles; 4) sequencing reads tend to have many errors due to technological infidelity.

To fill in this gap, we developed Monopogen, a computational framework that enables researchers to detect single nucleotide variants (SNVs) from a variety of single cell transcriptomic and epigenomic sequencing data. Monopogen leverages linkage disequilibrium (LD) data from an external reference panel to increase SNV detection sensitivity and genotyping accuracy. To detect novel SNVs that do not exist in the external panel, Monopogen uses Monovar [17], a probabilistic SNV caller that effectively accounts for allelic dropout and false-positive errors in single-cell sequencing data to identify rare germline SNVs, somatic SNVs or RNA editing events at the level of cell types or cell states. We demonstrated Monopogen as a novel computational framework that brings together population genetics and single cell data analytics to uncover genetic drivers of cellular-quantitative traits in ongoing single-cell-based studies.

## Results

### Workflow of Monopogen

Monopogen starts from individual bam files of single cell sequencing data, produced by single-cell RNA-seq (scRNA-seq), single-nucleus RNA-seq (snRNA-seq), single-nucleus Assay for Transposase-Accessible Chromatin using sequencing (snATAC-seq), etc (**Fig. 1a**). Sequencing reads with high alignment mismatches (default 4 mismatches) are removed. Putative single nucleotide variants (SNVs) are detected from pooled (across cells) read alignment wherever an alternative allele is found in at least one read. For SNVs that are present in an external haplotype reference panel, such as the 1,000 Genomes phase 3 (1KG3) panel, the input genotype likelihoods estimated by Samtools are further refined by leveraging linkage disequilibrium (LD) from the reference panel to account for genotyping uncertainty in sparse sequencing data. The loci showing persistent discordance after LD refinement are used to estimate a sequencing error model for de novo SNV calling (**Fig. 1b**). For remaining loci satisfying minimal total sequencing depth and alternative allele frequency cutoffs, Monovar [17] is then used to call novel SNVs at cell cluster level (Details shown in **Fig. S1a**). These loci can be further classified as germline, somatic, or RNA editing events based on whether they are detected in specific or all the clusters (**Fig. 1c**). The germline SNVs from **Fig. 1b** can be used for global or local ancestry inference (**Fig. 1d**), or cellular quantitative trait mapping when sample size is sufficient (**Fig. 1e**)

**Fig.1.**
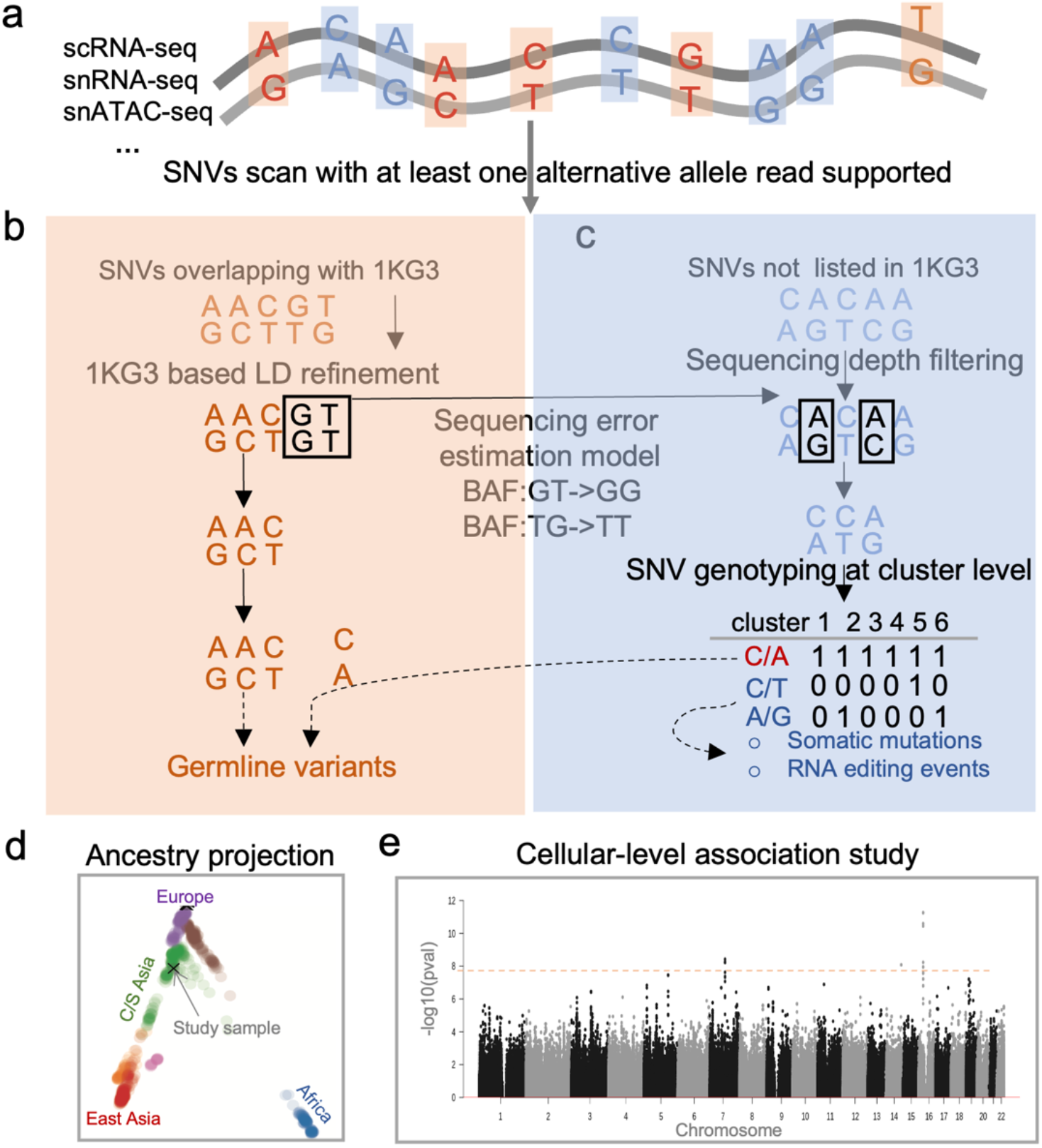
Monopogen workflow. **a,** Monopogen starts from individual bam files produced by single cell sequencing technologies including scRNA-seq, snRNA-seq, snATAC-seq, etc. Sequencing reads with multiple alignment mismatches (default 4) are removed. Putative SNVs are identified sensitively from pooled pileup containing at least one non-reference read. **b,** For SNVs present in the external reference panel (such as 1000 Genome 3; 1KG3), genotype likelihoods are further refined based on linkage disequilibrium (LD) in the reference panel. The loci showing persistent discordance are used to estimate a sequencing error model. BAF: B-allele frequency. **c,** For the remaining loci, Monovar is then used to call novel SNVs at cluster level (details in **Fig. S1a**), if there are sufficient sequencing depth and alternative allele frequency (calibrated by a sequencing error model). These loci can be further classified as germline, somatic, or RNA editing events based on whether they are detected in specific or all the clusters. **d,** Projection of study samples onto human genetic diversity panel (HGDP) enables genetic ancestry inference. **E,** Genome wide association study of cellular quantitative traits can be performed when there is sufficient sample size.

### Benchmarking of Monopogen performance on SNV calling

We used two single cell sequencing datasets (snRNA-seq from 4 retina tissue samples, sci-ATAC-seq from 2 colon tissue samples) having matched WGS data to evaluate SNV calling performance. In all these samples, the overall accuracy (**Methods**) of the Monopogen calls was higher than 95% for the germline SNVs present in the 1KG3 panel, 97% for **5/6** of the samples (**Fig. 2a**; **Table S1**). The high accuracy is largely due to the LD-based genotyping refinement. The overall accuracy without LD-based refinement, such as calls from Samtools and GATK was less than 73% (**Table S2**). Further examination shows that over 85% of the genotyping errors was misclassifying 0/1 as 1/1 (**Table S1**), due partly to allele drop artifacts in the single cell data.

**Fig.2.**
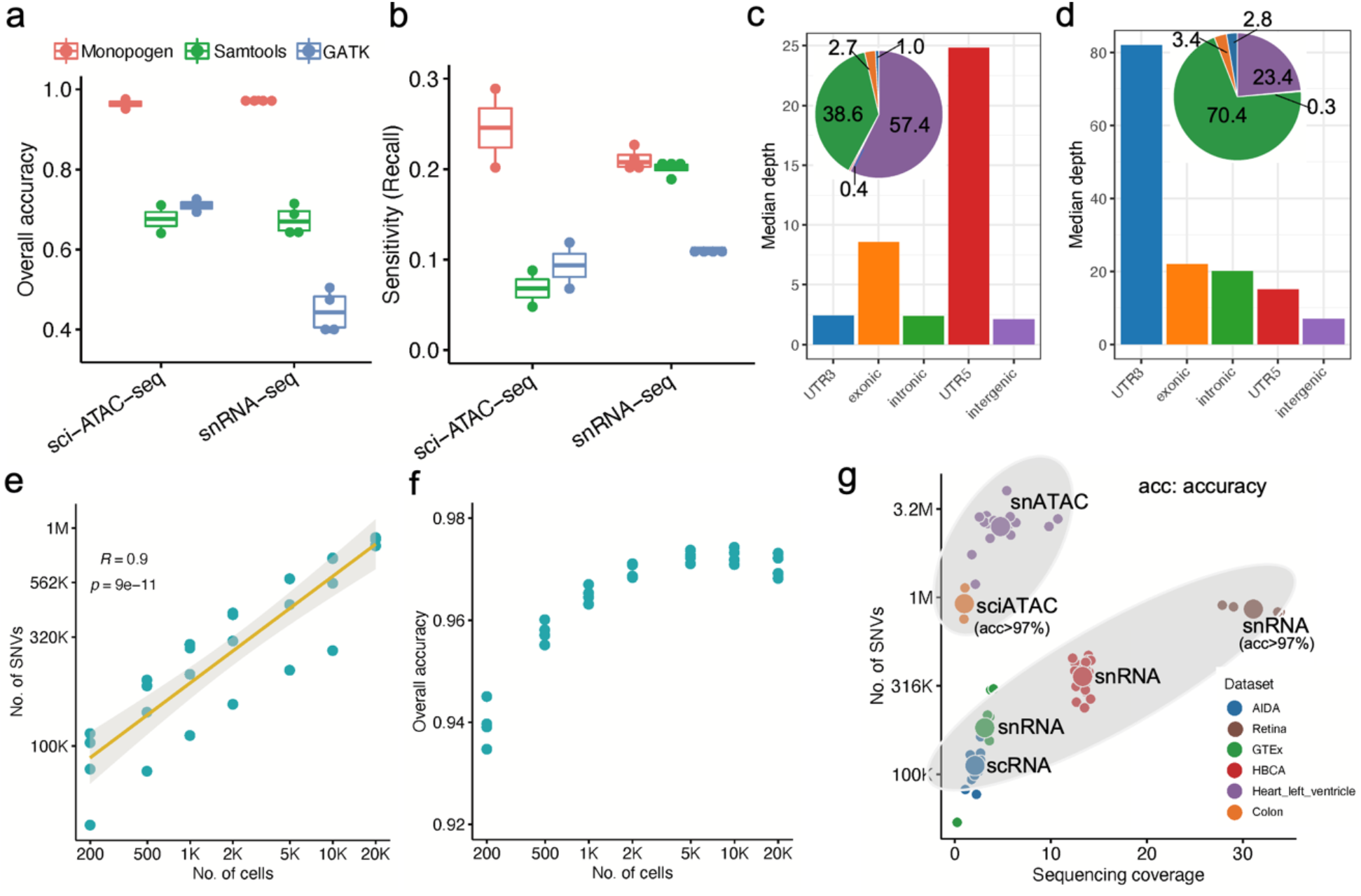
Benchmarking of Monopogen performance in various single cell sequencing platforms. **a**, Overall accuracy in representative snRNA-seq (4 samples) and sci-ATAC-seq data (2 samples) using matched WGS data as the gold standard, comparing Monopogen against Samtools and GATK. Each dot denotes one sample, bars denote means and boxes 95% standard deviations. **b.** SNV detection sensitivity (recall). Note, in **a-b** for Monopogen evaluation, only the SNVs present in the 1KG3 were considered. **c-d,** Median sequencing depth of SNVs found from sci-ATAC-seq data (**c**) and snRNA-seq data (**d**) over gene annotations. The pie charts show the percentage of SNVs in each category. **e,** Number of SNVs versus number of cells in the retina data via down-sampling. The X-axis and Y-axis are in logarithmic scale. Pearson’s correlations were applied to calculate the R and the p values. **f,** Overall accuracy versus cell number. **g**. Number of SNVs detected from six single cell sequencing datasets. The sequencing coverage was calculated as the *L* × *N*/(3.2 × 10^9^), where *L* is the read length, *N* the total number of reads in one sample. Each dot denotes one sample colored by dataset. The top ellipse covers samples from single-cell ATAC-seq data and the bottom ellipse samples from single-cell RNA-seq data.

In the retina snRNA-seq data, Monopogen detected 827K~905K germline SNVs, achieving a recall of 21% (**Fig. 2b; Table S1**). GATK and Samtools achieved a recall of 11%~20% at the expense of lower accuracy (<73%). Most (70.4%) SNVs from Monopogen were detected in intronic regions, only less than 7% in exonic regions (**Fig. 2d**). As expected, sequencing depth was substantially higher in genes than in intergenic regions. Off-target reads appear sufficiently leveraged to derive accurate genotypes through LD-based refinement.

In addition, Monopogen detected ~100K novel SNVs in the retina snRNA-seq data that are not presented in the 1KG3 panel, after performing sequencing depth filtering (>100) and sequencing error model calibration. The overall accuracy of this set is 35% and is 86% for the subset detected in more than 90% of the transcriptomic clusters determined by Seurat [18] (**Fig. S1 b; Table S3**).

In the colon sci-ATAC-seq data, Monopogen detected 752K~1.1M germline SNVs, achieving a recall of 25%. In contrast, the recall for Samtools and GATK were less than 12%. Most (57.4%) of the SNVs from Monopogen were found in intergenic regions and 38.6% in gene regions (**Fig. 2c**).

To evaluate Monopogen’s performance on sparser sequencing data, we ran Monopogen on down-sampled retina snRNA-seq data containing random subsets of 200 to 20,000 cells (~29.4K reads per cell; **Table S1**). We observed a linear relationship between the number of SNVs and cell numbers in logarithmic scale (**Fig. 2e**, Pearson correlation coefficient is 0.9). Monopogen detected ~100K SNVs from only 200 cells and 500K SNVs from 1,000 cells (**Fig. 2e**). Remarkably, despite of downsampling, the overall accuracy of Monopogen remained robust to cell number and was always higher than 94% (**Fig. 2f**).

We further evaluated Monopogen in four other cohorts: human breast cell atlas (HBCA; 20 donor samples), peripheral blood mononuclear cells from Asian Immune Diversity Atlas (AIDA; 20 donor samples), genotype-tissue expression project (GTEx; 7 donor samples), and human heart left ventricle atlas (65 samples). These datasets have a variety of cell number, number of reads per cell, and read length (**Table S4**). To make a fair comparison across datasets, we investigated the relationship between sequencing coverage and number of SNVs. As expected, Monopogen detected more SNVs from single cell epigenomics sequencing data than from single-cell transcriptomics sequencing data (**Fig. 2g**).

### Accurate global and local ancestry inference from single cell sequencing data

We performed genetic ancestry inference using genotypes called from Monopogen.

We projected via principal component analysis (PCA) the Monopogen-called snRNA-seq genotypes and the matched WGS genotypes of the 4 retina samples respectively onto a map, consisting of source samples with east Asia, America, Middle East, Europe, Oceania, Africa and central/south (C/S) Asia in the Human Genome Diversity Project (HGDP) [19]. We found that the PC coordinates were highly consistent between the WGS genotypes and the single-cell genotypes called by Monopogen (**Fig. 3 a-b)**. The mapping results were consistent with self-reported ethnicities for all the samples, including 3 Europeans and a self-reported Hispanic sample. We further performed local ancestry inference using RFMix [20]. On all the samples, the chromosomal painting results based on single cell data (**Fig. 3c-f; Fig. S3**) appeared highly consistent with self-reported ethnicities and with those obtained from the WGS data. For example, the source consistency across genomic bins was as high as 0.96 for one of the European samples (19D013) (**Fig. 3g**) and 0.90 for the Hispanic sample (19D015) (**Fig. 3h**). We did observe some genomic bins showing discrepant sources, due largely to sparseness of single cell derived SNVs in those regions. Remarkably, the global ancestry inference results remained largely unchanged when downsampling the data to only 200 cells (~29.4K reads per cell) (**Fig. S4**).

**Fig.3.**
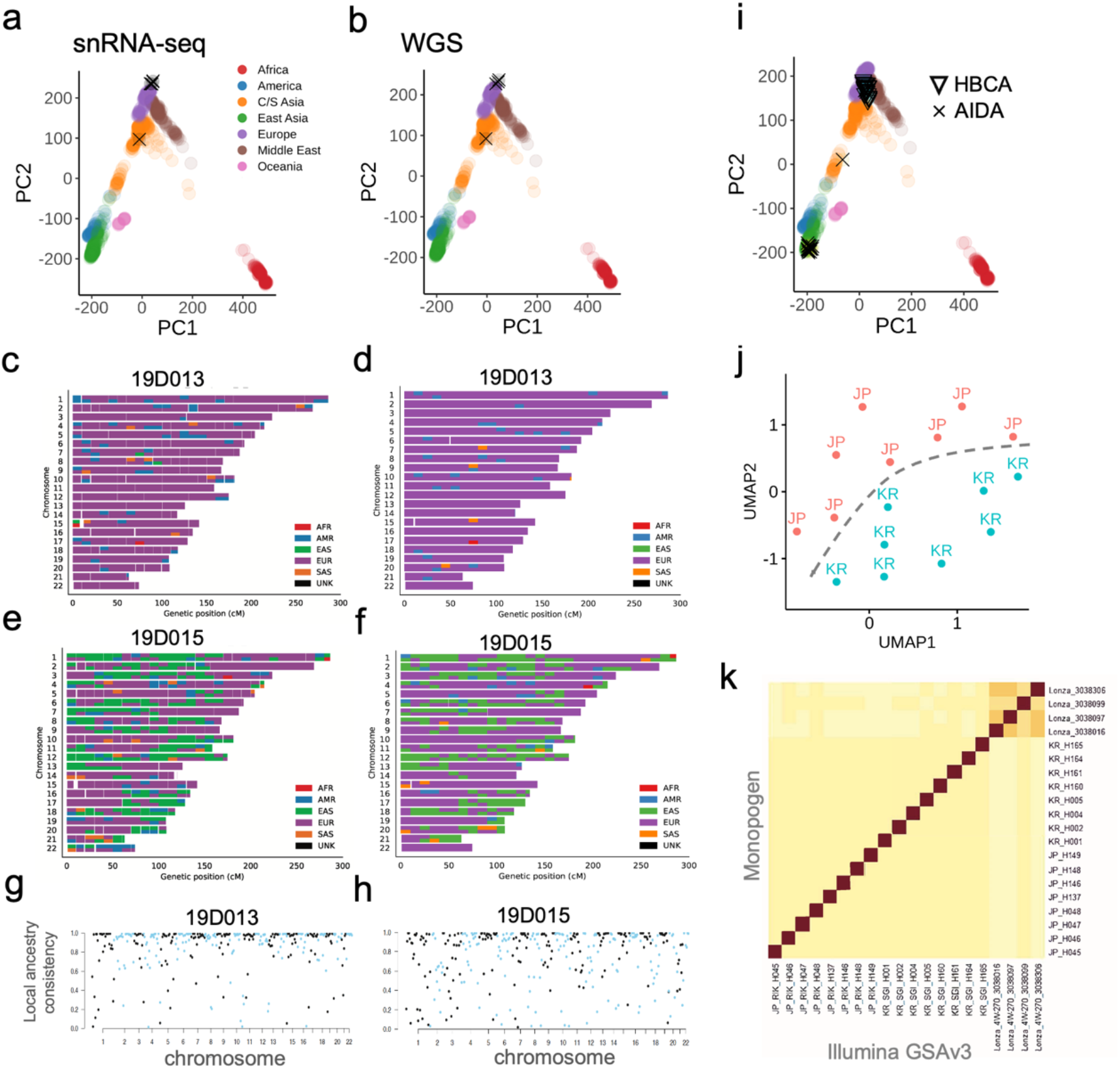
Global and local ancestry inference using single-cell genotypes derived by Monopogen. **a-b,** Genetic ancestry of the 4 retina samples using Monopogen genotypes derived from snRNA-seq data (**a**) and genotypes from matched WGS data (**b**). Colored dots represent 897 individuals in the HGDP reference panel and black crosses the retina samples. **c-d.** Local ancestry inference of an European sample 19D013 using genotypes from the snRNA-seq (**c**) and the WGS (**d**) data. The 3202 phased genotypes from 1KG3 were used as the reference for local ancestry inference. Colors in each chromosome denotes the inferred source ancestry with a bin size of 1 cM. **e-f**, Local ancestry results from an admixed sample 19D015. **g-h,** Local ancestry inference accuracy for 19D013 (**g**, overall score: 0.96) and 19D015 (**h**, overall score: 0.90). Each dot denotes the ancestry accuracy for each segment (1cM). **i**, PCA-projection analysis shows ancestry of samples in the AIDA and the HBCA cohorts. HBCA: Human Normal Breast Cell Atlas; AIDA: Asian Immune Diversity Atlas. **j.** UMAP of Korean and Japanese samples in the AIDA using genotypes called by Monopogen. **k**. Concordance between Illumina GSAv3 genotyping array data and Monopogen calls across the AIDA samples. Darker colors denote a higher level of concordance between two data modalities. Calculation of the concordance scores is detailed in **Methods**.

We also performed projection analysis on another 40 samples in the HBCA and the AIDA cohorts which do not have matched WGS data. Again, the global ancestry inferred from single cell sequencing were consistent with self-reported ethnicities except for one putative admixed sample in the AIDA cohort (**Fig. 3i**). In the AIDA cohort, it is difficult to separate Japanese and Korean samples by PCA-projecting them onto the HGDP panel. However, these two populations can be well separated by performing independent UMAP analysis using Monopogen derived genotypes (**Fig. 3j**). Furthermore, Monopogen shows consistent performance in identifying donor-specific SNVs in the AIDA samples, based on the concordance of Monopogen-derived genotypes and Illumina GSAv3 genotypes (**Fig. 3k**), demonstrating the possibility of distinguishing individuals from the same ancestry. This indicates that the LD-based genotyping refinement from commonly used 1KG3 panel did not over-correct genotypes on sub-population or individual level, despite sparse sequencing coverage.

### Genome-wide association study of cellular quantitative traits

Genome-wide association studies (GWAS) have implicated causal variants in non-coding, regulatory regions. However, establishing causality of these variants remain challenging, even with additional results from quantitative traits mapping such as expression quantitative trait loci (eQTLs), methylation quantitative trait loci (meQTLs), etc. That may be due to hidden genetic effect variations across cell types or cell states (such as cell differentiation or activation). Previous bulk-based data disguised these relations. It is possible to uncover these hidden relations by associating, in a cell-type specific manner, genotypes inferred by Monopogen and quantitative traits estimated from single cell sequencing.

Characterizing genetic contribution to metabolic processes in healthy cardiac, such as ATP production, is necessary to understand metabolic alterations in heart failure (HF) and heart hypertrophy [21]. Epigenetic mechanisms and transcription factor networks are also essential for differentiation of cardiomyocytes [22]. However, it remains largely unknown how genetics affect the metabolism and the epigenome of cardiomyocytes.

As a demonstration, we collected snRNA-seq and snATAC-seq data of ~4M cells generated from human heart left ventricle tissue samples of 65 donors, 54 of which have data from both modalities. Around 791K SNVs in snRNA-seq and 2.59M SNVs in snATAC-seq were identified from Monopogen (**Table S5**). The variant calling consistency between two modalities was as high as 97% at overlapping loci (**Fig. S5 b-c**). Variant calls were further merged for samples of paired modalities.

Ancestry admixture analysis using inferred genotypes shows that this cohort contains samples with diverse ancestry: European (71.1%), Asian (10.2%) and African (8.5%). Six samples appeared admixed (**Fig. S6a**).

To explore the cardiac metabolism process, we extracted cardiomyocyte cells from each sample by annotating cells using the human heart Azimuth database (**Fig. 4b; Fig.S6b**). Using pathway expression level as a proxy for ATP metabolism level, we derived cardiac ATP metabolism level by aggregating the expression levels of 216 genes in *GO_ATP_METBOLIC* pathway (see **Methods**). We performed association analysis using the GCTA tool [23], including the top five ancestry PCs as covariates. P value of 10^-5^ was used as the threshold to identify potential associations due to the small sample size. The inflation factor of Quantile-Quantile plot was close to 1 (0.983, **Fig. S7a**). A total of 250 variants were associated with cardiac ATP metabolism score (*p*-value<10^-5^), which can be further binned into 42 gene regions (**Table S6**), including 5 genes (at least two variants supported) with p-value<10^-6^ (**Fig. 4c**). Among genes in the regions, *IGFBP3* and *FBXL22* are well known to affect adult cardiac progenitor cells [24] or cardiac contractile function [25]. *ADO* functions as an oxygen sensor involved in N-degron pathways [26]. These associations further confirm the tight coupling of ATP production and myocardial contraction, which is essential for normal cardiac function [27]. *AGAP1*, indicated by its tag SNV (rs6714660) (**Fig. 4d**), is involved in cardiac ATP production in Krebs cycle [21].

**Fig.4.**
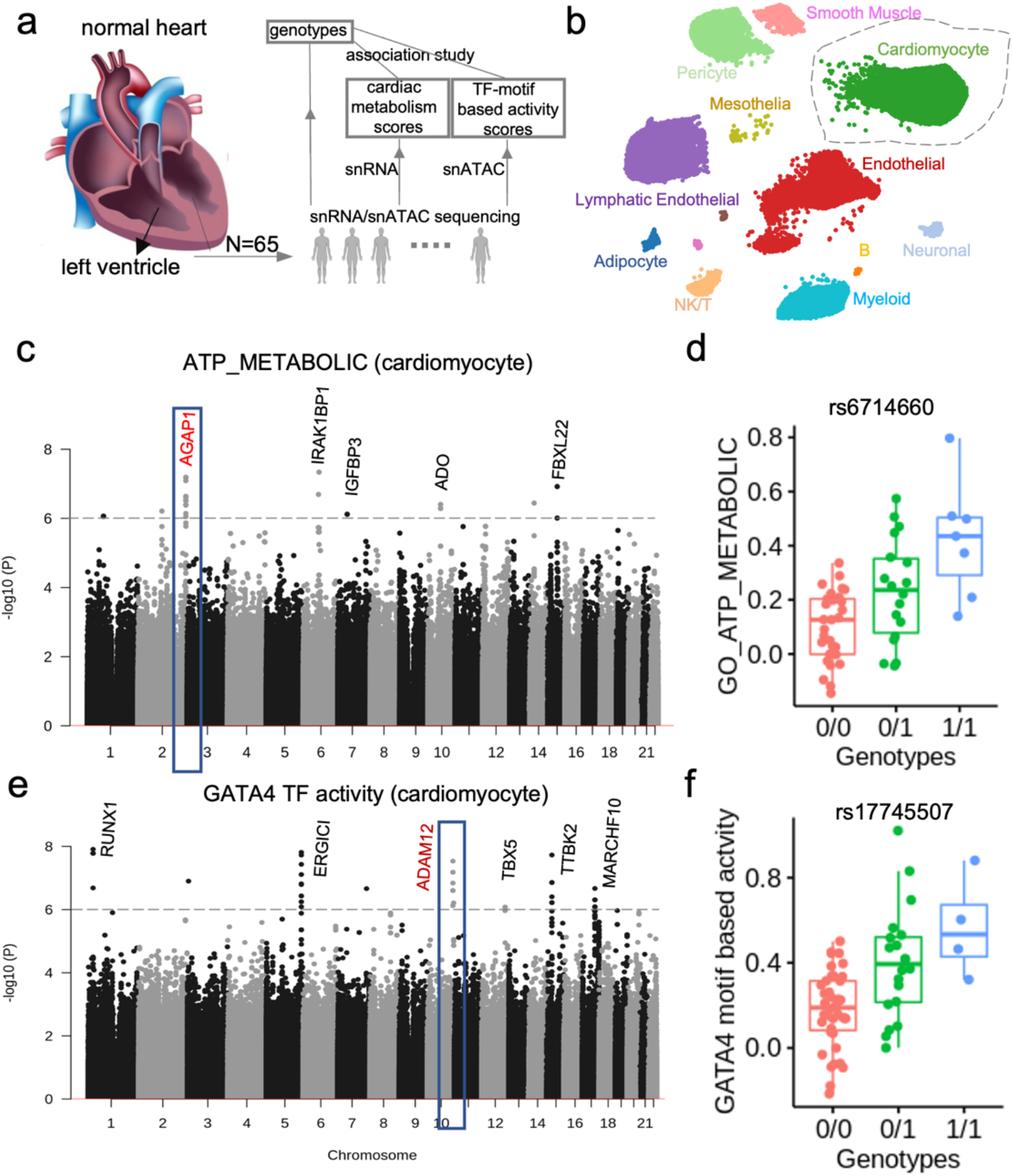
Genetic association study of cardiomyocyte 331 molecular traits using snRNA-seq and snATAC-seq data from heart left ventricle tissues. **a,** Analysis workflow. Details can be seen in **Methods**. **b**, A UMAP of snRNA-seq cells colored by cell types annotated using Azimuth heart database. **c,** Manhattan plot showing association of Monopogen SNVs with pathway scores of ATP_METABOLIC in cardiomyocytes. The gray line denotes the p-value threshold of 10^-6^. Genes closest to the top scoring loci are labeled. **d**, Boxplot shows the difference of ATP_METABOLIC scores across the three genotypes of rs6714660 (one of the leading variants in *AGAP1*). **e,** Manhattan plot showing association of SNVs with the *GATA4* motif-based transcription factor activity level in cardiomyocytes. The gray line denotes the p-value threshold 10^-6^. **f**, Boxplot shows the difference of *GATA4* activity level across the three genotypes of rs17745507 (one of the leading variants in *ADAM12*).

We also derived transcription factor (TF) activity scores from the snATAC-seq data (see **Methods**). We then scanned for genetic variants associated with the activity level of *GATA4*, one of the most important TFs highly activated in cardiomyocytes at various developmental stages. The inflation factor of Quantile-Quantile plot was close to 1 (0.984, **Fig. S7b**). A total of 257 variants were identified (*p*-value<10^-5^), which can be further binned into 42 gene regions (**Table S7**), 6 of which (at least two variants supported) with p-value<10^-6^ (**Fig. 4e**). Among the genes in the regions, *TBX5-GATA4* and *RUNX1-GATA2* complexes are well known for their interdependence in coordinating cardiogenesis [28–31]. A particularly interesting gene *ADAM12*, indicated by its tag SNV (rs17745507) (**Fig. 4f**), is known to play an important role in cardiac hypertrophy by blocking shedding of heparin-binding epidermal growth factor (HB-EGF) [32]. These results indicate a potential novel association between *GATA4* and cardiac hypertrophy through the mediation of *ADAM12*. Also identified were some variants (p-value <10^-5^), locating in zinc-finger family gene *such as ZNF595* and *ZNF750* which are acting as cofactors with the zinc finger transcription factor *GATA4* (**Table S7**).

In summary, we were able to, for the first time with the aid of Monopogen, reveal potential genetic determinants of cardiac health via metabolic and epigenomic trait mapping of cardiomyocytes, despite the relatively small sample size. Associations identified in this fashion may lead to better understanding on the pathogenicity of non-coding variants in a cell-type aware manner.

## Discussions

In this study, we developed Monopogen, a computational tool enabling researchers to identify SNVs at high accuracy from sparse single cell transcriptomic and epigenomic sequencing data. Single cell sequencing technologies, like other targeted sequencing technologies [33–35], can generate reads that map outside of the target regions, which has become a rich, under-utilized resource for genomic variant discovery. By leveraging these reads, in conjunction with the known LD patterns in major human populations, Monopogen identified around 100K~1M SNVs in 10X Chromium single cell or nucleus RNA-seq data, and 1M~2.5M SNVs in single cell ATAC-seq data at genotyping accuracies higher than 0.95. We found through downsampling experiments that Monopogen can be applied in most single cell sequencing datasets, including those with low (~200) cell numbers. Although not evaluated in this work, there should be no barrier to apply Monopogen on data produced by other single-cell sequencing platforms such as the full-length smart-seq [36] and single cell DNA sequencing [37, 38]. With SNVs called by Monopogen, global and local ancestry inference can be reliably performed in studies that have only single-cell sequencing but not bulk sequencing or array-based genotyping data, which greatly increases the chance of discovering novel genetic factors underlying diverse cellular quantitative traits and disease.

Health disparity is a significant socioeconomic challenge. Ongoing large-scale single-cell studies (such as HCA and the CZI genetic ancestry network) are aiming at creating a genetically unbiased reference and avoiding the Eurocentric biases in previous human genetic studies [39]. Our study has clearly shown that single-cell sequencing data can potentially be utilized as a resource to not only determine the genetic ancestry of study samples but also expand the reference to further delineate human populations. For example, we found a clear separation of Japanese and Korean samples in the AIDA cohort based on variants and genotypes determined from single-cell data by Monopogen. Moreover, although our analysis and assessment were based on publicly available reference population databases such as 1KG3, we expect that the power of variant calling and ancestry inference will become greater when using local population panels [40, 41] or proprietary databases with larger population size and greater diversity.

Monopogen adds a genomic modality to current single-cell transcriptomic and epigenomic assays [9, 42, 43], which makes it possible to use these assays for functional genetics investigations. For example, we identified novel SNVs that are associated with the metabolism and epigenetic regulation of cardiomyocytes in heart samples. Many similar analyses can be performed, for example, identifying genetic determinants of cancer immune response using pan-cancer single cell T cell atlas data [44].

Monopogen is efficiently implemented in Python. It starts from aligned BAM files and adopts the framework implemented in GotCloud [45] for Trans-Omics for Precision Medicine Program (TOPMed), by automatically splitting the genome into small overlapping chunks (defined by the users), performing variant scan and LD refinement in massive parallelization for individual chunks and merging the results. For individual sample with each chromosome as one chunk, the average CPU time is 1.5h for one chunk with maximal memory usage less than 11 Gb. It also supports population-scale single cell sample genotyping. With 10Mb region as one chunk, joint calling of 65 snATAC-seq samples takes ~6.2h CPU times for each chunk with maximal memory usage less than 10Gb.

Our study has several limitations. Although Monopogen can potentially detect all types of SNVs, we focused on germline SNVs in the reference panels. The putative novel variants called by Monopogen are likely important for many other investigations, such as performing clonal lineage tracing in cancer evolution studies [17, 38, 46]. However, due to challenges in experimental validation, the accuracy of these novel variants was not fully assessed in this study and will require further investigation. We did observe reasonable accuracy (>35%) of *de novo* SNV calling in the retina snRNA-seq data, supporting the use of Monovar, error-filtering rules and choice of parameters. In the human heart left ventricle analysis, we demonstrated the utilization of Monopogen-called genotypes to identify associations of ATP metabolism and *GATA4* activity levels in one cell type, cardiomyocytes. In the context of discovery, such analysis can be extended to other cell types and other cellular quantitative traits of interest that could be objectively measured. However, such association analysis should be guided by strong prior knowledge to reduce the burden of multiple hypothesis testing.

In summary, we developed a computational tool Monopogen to enable population genetic investigations on single cell sequencing data, which can lead to immediate benefits on genetic ancestry mapping and association analysis using current large-scale single cell atlas data [10, 11]. In long term, with the increasing generation of sparse single cell sequencing data and expansion of data modalities, our work will become increasingly relevant for assessing effects of genetic ancestry and discovering genetic mechanisms underlying complex traits in human populations and diseases.

## Methods

### 1. Monpogen workflow

#### Reads filtering

Monopgen starts from individual bam files of single cell sequencing data including scRNA-seq, snRNA-seq or snATAC-seq etc. Sequencing reads with high alignment mismatches (default 4 mismatches) and lower mapping quality (default 20) are removed.

#### SNV discovering

We first scan the putative SNVs in a sensitivity way. Any loci are detected from pooled (across cells) read alignment from one sample wherever an alternative allele is found in at least one read. We do not perform genotyping at this step since the sequencing depth is low for a majority of candidate SNVs. For each candidate SNV locus *m* with observed sequencing data information *d*, we record its genotype likelihoods (GL) that incorporate errors from base calling and alignment as

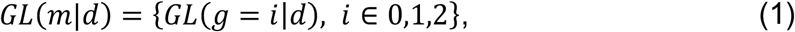

in which *g* = 0 denotes homozygous reference allele, *g* = 1 denotes heterozygous, and *g* = 2 denotes homozygous alternative allele. Calculation of *GL*(*g*|*d*) is performed using Samtools *mpileup* tool [14].

#### Germline variant calling refinement

Given single cell RNA sequencing data has high genotyping uncertainties and is quite sparse, we leverage the linkage disequilibrium (LD) from the 1000 Genome 3 (1KG3) database to further refine the genotype likelihoods. The 1KG3 database was downloaded, which includes 3,202 samples with a total of ~80M SNVs after quality control and genotypes phased using Shapeit2,

Focusing on putative SNVs existing in both the 1KG3 panel and the single cell sequencing data, we combine multiple consecutive putative SNVs within a large interval of 0.5cM as a block. Denotes *M* the list of SNVs blocks in chromosome order and *H* the set of reference haplotypes (|*H*| = 6,404). The Beagle hidden Markov model (HMM) [47, 48] is used to identify the target haplotype of *M* SNV blocks from sparse sequencing data. The HMM includes four parts: 1) definition of state space; 2) initial probabilities, 3) transmission probabilities; and 4) emission probabilities. Briefly,

1. Denote the model states at a SNV block *m* ∈ *M* as *H_m_* = {(*m, h*): *h* ∈ *H*}.
2. For the first SNV block in *M* (with chromosome order), an initial probability of *1/H* is assigned to *H*_1_.
3. Denote the random variable *S_m_* ∈ *H_m_* as the state of the HMM and its transition probabilities:

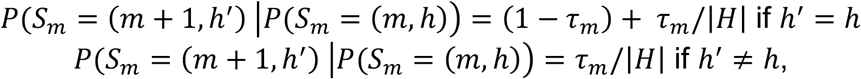

in which *τ_m_* = 1 – *e*^-*ρ_m_*/|*H*^ is the probability of transitioning to a random state at the next block, *p_m_* = 4 *N_e_ r_m_, N_e_* the effective population size, and *r_m_* the genetic distance between block *m* and *m* + 1.
4. In terms of emission probabilities, suppose that the SNV block *m* ∈ *M* consists of *m_k_* constituent genotyped markers and has *m_k_* distinct allele sequences in the reference and single cell sequencing sample. The state (*m, h*) emits the allele sequencing present on haplotype *h* with probability max (1 – *m_l_ε*, 0.5), and it emits each of other (*k* – 1) segregating allele sequences with probability min(*m_l_ ε*, 0.5) / (*k* – 1). *ε* is the allele error rate setting with 0.0001.

Based on above designed HMM space, the forward-backward algorithm in Beagle is used to estimate *P*(*S_m_* = (*m, h*)), conditioning on the observed alleles on each target haplotype. Note that the reference panel SNVs absent in the study sample are not included in the process.

Equation (1) is further updated as the genotype probabilities conditioning on the haplotypes in the reference panel as

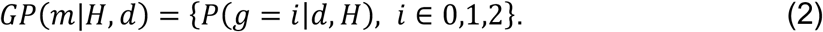

#### Sequencing error modeling

For each locus *m*, we calculate the observed genotype as the one with the highest posterior probability from equation (1) and (2), respectively. Denote

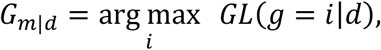

and

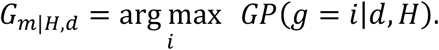

The final genotype of locus *m* is set as *G*_*m*|*Hd*_ if *G*_*m*|*Hd*_ = *G*_*m*|*d*_.

The heterozygous loci that are imputed to homozygotes are considered as sequencing errors (i.e., *G*_*m*|*H,d*_ = 0 and *G*_*m*|*d*_ = 1, 2). We classify these discordance into 12 categories:

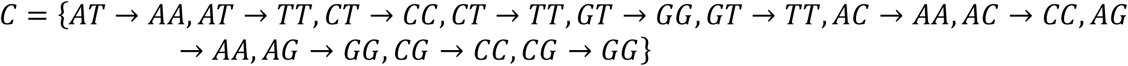

The median B-allele frequency (BAF) across all inconsistent loci in each category *c* is denoted as *BAF_C_*. This is considered as the threshold to separate the sequencing error from the true heterozygous. SNVs with *G*_*m*|*H,d*_ = *G*_*m*|*d*_ are retained as the germline SNVs (i.e., SNVs). Others are only used to build the sequencing error model and are not included in the final genotyping call set.

#### De novo SNV calling

For putative SNVs absent in the 1KG3, we implement two filters: 1) the total sequencing depth filtering (default 100); 2) BAF less than the threshold from above sequencing error model. For example, one putative SNV genotyped as *A/T* with its BAF lower than max {*BAF*_*AT*→*AA*_, *BAF*_*AT*→*TT*_} is removed due to difficulties in separating true heterozygotes from sequencing errors.

Monovar [17] is then used to perform SNV calling and genotyping on remaining loci at cluster or cell type level. Briefly, cell cluster identification can be obtained either by clustering on single cell profiles or using reference-based cell type annotation [18]. To reduce the computational time, only reads covering these candidate loci are extracted and then split into different bam files based on their cluster identifies (**Fig. S1a**). Monovar can be run on these bam files (each is one cluster or cell type) to detect putative SNVs, which can be further classified as private germline SNVs, somatic SNVs, RNA editing events, or unknown errors based on whether they show cluster specific mutational pattern.

### 2. Genotyping calling evaluation

Six single cell samples in our study have matched WGS data that were treated as the gold standard. For each sample, only bi-allelic loci having at least one alternative alleles (i.e., genotype is 0/1 or 1/1) were extracted from the two call sets, denoting as *N* (Monopogen-called) and W(WGS-called). Variants from the WGS data were called using GATK. The sensitivity (recall) was defined as |*N* ∩ *W*|/|*W*| and specificity (precision) as 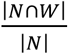. The genotyping accuracy was defined as the fraction of identical genotypes in the |*N* ∩ *W*| overlapping SNVs. The overall accuracy was defined as the specificity multiplied by the genotype accuracy.

The genotype concordance of the Monopogen-called genotype data versus the AIDA Illumina GSAv3 genotype data was computed by first counting the number of matching alleles between the Monopogen and the Illumina GSAv3 results for loci found in both sets. The minimum possible concordance score per Monopogen calls (accounting for some match always being possible in the case of heterozygous genotypes) was subtracted, and the resulting scores were then normalized against the number of loci evaluated.

We also randomly selected 3 samples from 65 left ventricle samples having paired snRNA-seq and snATAC-seq data. The genotyping accuracy was estimated based on SNVs detected in both modalities.

### 3. Global and Local ancestry analysis

#### PCA-projection analysis

To identify global ancestry of single cell sequencing samples, we downloaded genotypes from Human Genotyping Diversity Panel (HGDP), which includes 938 individuals (covering 53 populations worldwide) and 632,958 SNVs with MAF>1%. Denote ***R**_N×L_* as genotypes of the HGDP samples (*N* = 897, *L* number of SNVs), and *g*_1×*L*_ as the Monopgene-called genotype vectors from the single cell sequencing samples. The LASER (Trace module) [49] was used to project each sample to the HGDP. Briefly, the top *K* PCA coordinates of HGDP were calculated as *Y_N×K_* by applying eigen value decomposition (EVD) on the genetic relationship matrix ***RR**^T^*. Denote 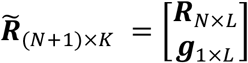, the top *K′* PCA coordinates (*K′* ≥ *K*) were calculated as 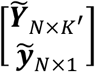 by applying EVD on 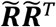.

In this case, there were two sets of coordinates for the same *N* reference individuals ***Y**_N×K_* and 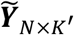. Projection procrusters analysis was used to find an orthonormal projection matrix ***A**_K′×K_* and an isotropic calling factor *ρ* such that 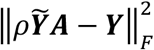 is minimized, where 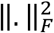 represents the square of Frobenius norm. Once ***A**_K′×K_* and *ρ* were solved, the sample specific PCA-projection coordinates on HGDP panel can be calculated as 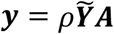. In this study, *K* and *K′* were set as 3 and 20, respectively. The PC coordinates of 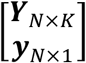 were used for PCA-projection visualization.

#### Fine-scale ancestry inference

The local ancestry components of single cell sequencing samples were calculated using RFMix tool [20] with the phased haplotypes from the 1000 Genomes 3 as reference source. Monopogen-called genotypes were first phased using Beagle, followed by inputting to the PopPhased version of RFMix v.1.5.471 [20] with the following flags: *-w 0.2, -e 1, -n 5, --use-reference-panels-in-EM, --forward-backward EM*. The node size of 5 was selected to reduce bias in random forests resulting from unbalanced reference panel sizes (AFR panel N = 504, EUR panel N = 503, and NAT panel N = 43). We set the segment size as 1cM and used EM iterations to improve the local ancestry calls without substantially increasing computational complexity. The RFMix output was collapsed into haploid bed files, and “UNK” or unknown ancestry was assigned where the posterior probability of a given ancestry was <0.90. These collapsed haploid tracts were used for local ancestry component visualization.

#### Evaluation of fine-scale ancestry inference

The RFMix tool was run on both single cell sequencing and WGS genotypes from matched samples with segment size being 1cM. For each segment, the ancestry component percentage for each source population was recorded. The local ancestry consistence index was calculated as the correlation of ancestry component vector between the two call sets.

### 4. GWAS on cellular quantitative traits

#### Variant calling on Human heart left ventricle samples

The dataset of heart left ventricle tissues was downloaded from ENCODE project. There are 54 donors sequenced with snRNA-seq and 65 with snATAC-seq, among which 54 are paired. For downstream association study, SNV calling of 54 snRNA-seq and 65 snATAC-seq samples were performed separately using Monopogen, followed by removing MAF<1%. Variant calls were further merged for samples of paired modalities (**Table S4**).

#### Cell type annotation on snRNA-seq profiles

We also downloaded the matched snRNA-seq gene expression profiles and performed a series of filtering to remove cells expressing lower than 200 and higher than 10,000 genes, and with mitochondrial gene percentage higher than 15%, using Seurat V4 [18].

Cell type annotation was performed by uploading all the cells of each sample to the online Azimuth heart database in Seurat V4 [18]. Cells with predicted cell type probability score lower than 0.9 were removed. Only cells annotated as cardiomyocytes were extracted for downstream association study.

#### Cell type annotation on snATAC-seq profiles

Starting from the fragment files of snATAC-seq samples, we used Signac pipeline [50] to re-call peaks in each sample and combine them into an unified set after removing peaks of width<20bp and > 10kb, leading to a total of 488,652 peaks. The gene-level chromatin accessibilities were derived using *geneActivity* module by aggregating peaks in gene promoters plus upstream 2kb. The cell type annotation was also performed using the online Azimuth heart database under the same quality control criteria as did in the snRNA-seq analysis.

#### Calculation of cellular quantitative traits

We used pathway expression level as a proxy for ATP metabolism level. We downloaded 216 genes from *GO_ATP_METBOLIC* pathway. We derived cardiac ATP metabolism level at single cell resolution by aggregating the expression levels of 197 genes (197/216) detected in the snRNA-seq data. The calculation was performed using *AddModuleScore* module in Seurat. In snATAC-seq, transcription factor *GATA4* motif-based activity was calculated for each cell using ChromVAR [51].

#### Association study

GCTA [23] was used to calculate a genetic relationship matrix (GRM) among single cell sequencing samples. The association studies on ATP metabolism level and GATA4 activity level were performed using its *fastGWA-mlm* option with the input of GRM and covariates as top five ancestry PCs. Only variants with MAF>10% were considered for association studies. The inflation factor of Quantile-Quantile plots was calculated using the R package *qqman* to examine whether there is population stratification in our genome-wide scan. Manhanttan plot was used to show the p-value across the whole genome with *pval*=10^-5^ as potential significant associations with cellular traits. The significant loci were further grouped into bins based on their closest genes. The nearest genes to significant loci were annotated.

### 5. Comparison with other SNV callers

For a fair comparison with Monopogen, both Samtools [14] and GATK [52] were run on bam files after the same filtering with Monopogen. For Samtools, the *mpileup* option was used to transform base calling and alignment information into the genotype likelihoods, followed by variant calling using Bcftools. The GATK was run using the *HaplotypeCaller* mode with default settings.

## Data availability

The sci-ATAC profiles from the two transverse colon samples were downloaded from ENCODE database at https://www.encodeproject.org/files/ENCFF354SCV/ and https://www.encodeproject.org/files/ENCFF491HQL/. The matched VCF files for WGS genotypes were from accession https://www.encodeproject.org/files/ENCFF944WLM/ and https://www.encodeproject.org/files/ENCFF907ASL/.

The snRNA-seq and snATAC-seq profiles from the human heart left ventricle tissues of 65 donors were downloaded from ENCODE database at https://www.encodeproject.org/matrix/?type=Experiment&assay_title=snATAC-seq&assay_title=scRNA-seq&biosample_ontology.term_name=heart+left+ventricle.

The 12 scRNA-seq samples with matched WGS genotypes were downloaded from GTEx database with https://anvil.terra.bio/#workspaces/anvil-datastorage/AnVIL_GTEx_V9_hg38.

The 1KG3 genotypes were downloaded from https://ftp.1000genomes.ebi.ac.uk/vol1/ftp/data_collections/1000G_2504_high_coverage/working/20201028_3202_phased/.

The HGDP panel genotypes were downloaded from http://csg.sph.umich.edu/chaolong/LASER/HGDP-938-632958.tar.gz.

The single cell profiles of 20 HBCA scRNA-seq samples, 20 AIDA samples, and 4 retina samples were generated as part of the cell atlas and genetic ancestry networks organized by the Chan Zuckerberg Initiative, which are available through the networks.

## Software Availability

Monopogen is available in open source at https://github.com/KChen-lab/Monopogen.

## Acknowledgements

This project has been made possible in part by Human Cell Atlas Seed Network Grant CZF2019-02425 and CZF2019-002432, Genetic Ancestry Network Grant CZF 2021-239847 from the Chan Zuckerberg Initiative DAF, an advised fund of Silicon Valley Community Foundation, grant U01CA247760 to KC and the Cancer Center Support Grant P30 CA016672 to PP from National Cancer Institute. This work is also supported by the CPRIT Single Cell Genomics Center (RP180684) gran to NN. We thank Norbert Tavares at CZI for advising the study, Joseph Powell and Drew Neavin for suggestions/discussions, Weiyi Xu from Baylor College of Medicine for suggestions on left ventricle single cell studies, Hamim Zafar and Luay Nakhleh for Monovar implementation/maintenance.

## Supplementary Figures

**Fig.S1.**
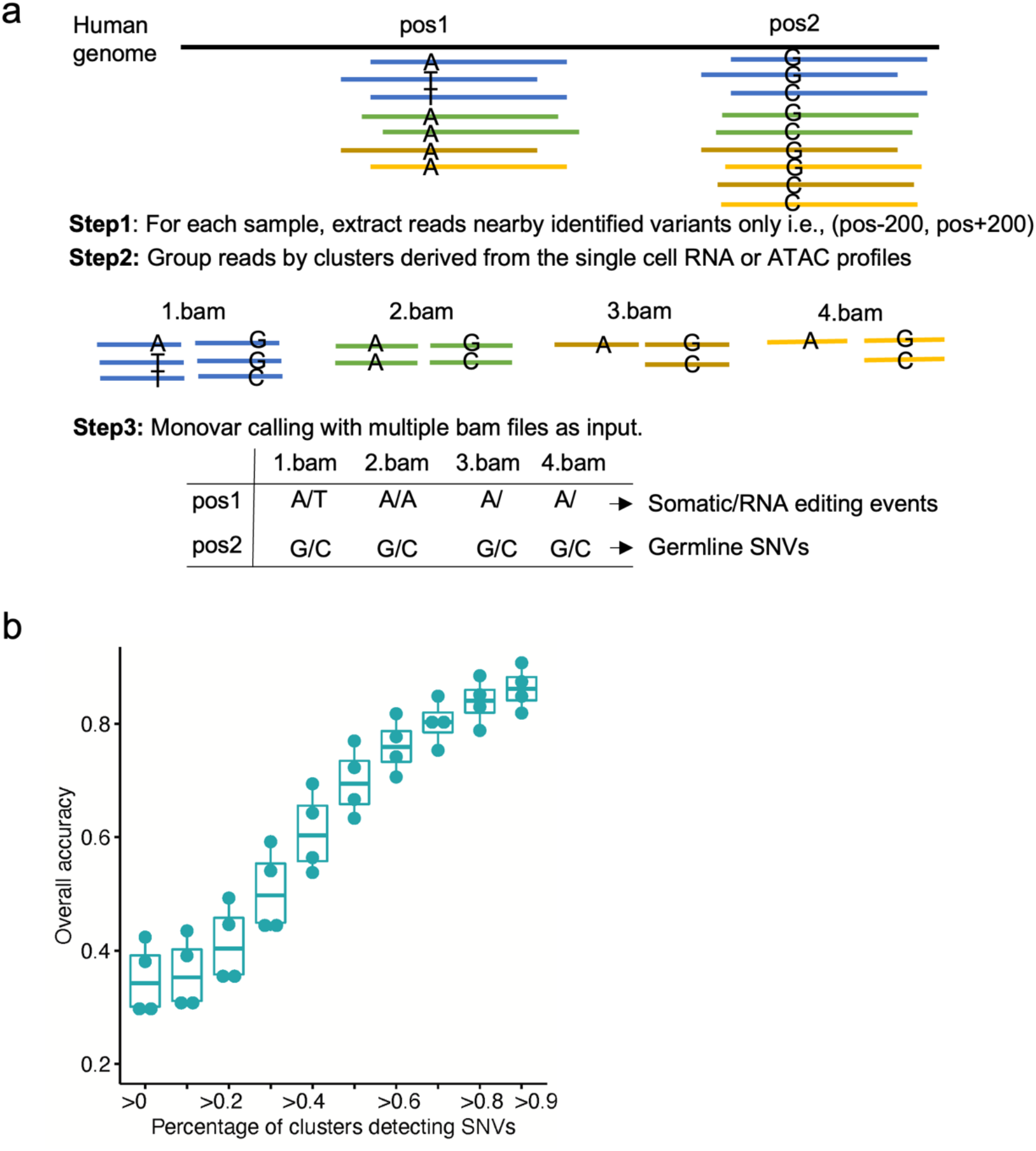
De novo SNV calling using cluster level information. **a,** De novo SNV calling of Monovar at the cluster level. Only reads covering putative SNV locations are extracted. They are then split into different groups based on their cluster identifies defined from expression level, followed by Monovar calling with multiple bam files as input (each bam file stores sequencing reads from one cluster). The germline SNVs or somatic/RNA editing events are further distinguished based on whether they are detected in specific or all the clusters. **b**, Overall accuracy of novel SNVs not present in the 1KG3 panel (Y-axis) with respect to the fraction of clusters in which the SNV was detected (X-axis). Each dot denotes one sample from retina studies.

**Fig.S2.**
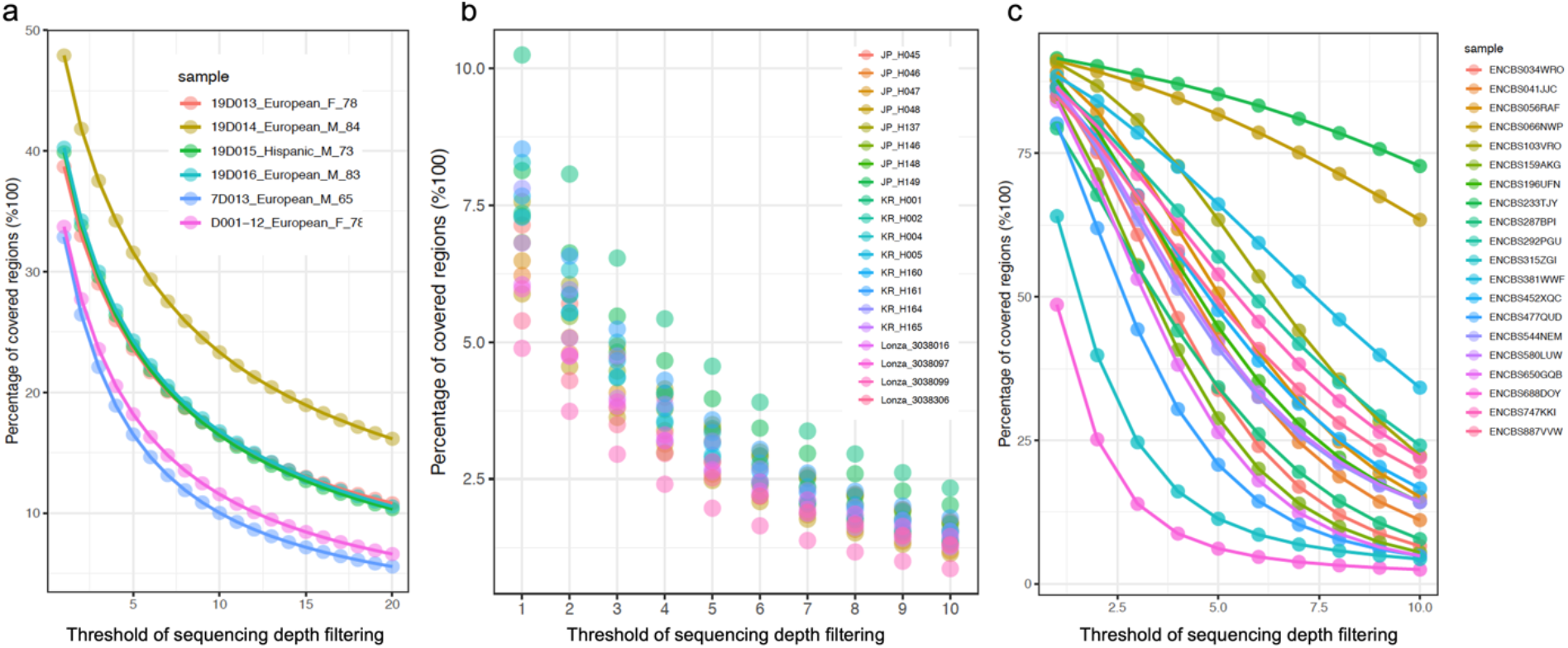
Sequencing depth survey on single cell sequencing datasets. **a**, Human retina snRNA-seq data (6 samples).**b,** AIDA scRNA-seq data (20 samples), and **c** heart left ventricle snATAC-seq data (randomly select 20 samples from 65 donors), respectively. Each curve denotes results for one sample. The X-axis denotes the sequencing coverage threshold and Y-axis denotes the percentage of genome region covered by more than the specified depth threshold.

**Fig S3.**
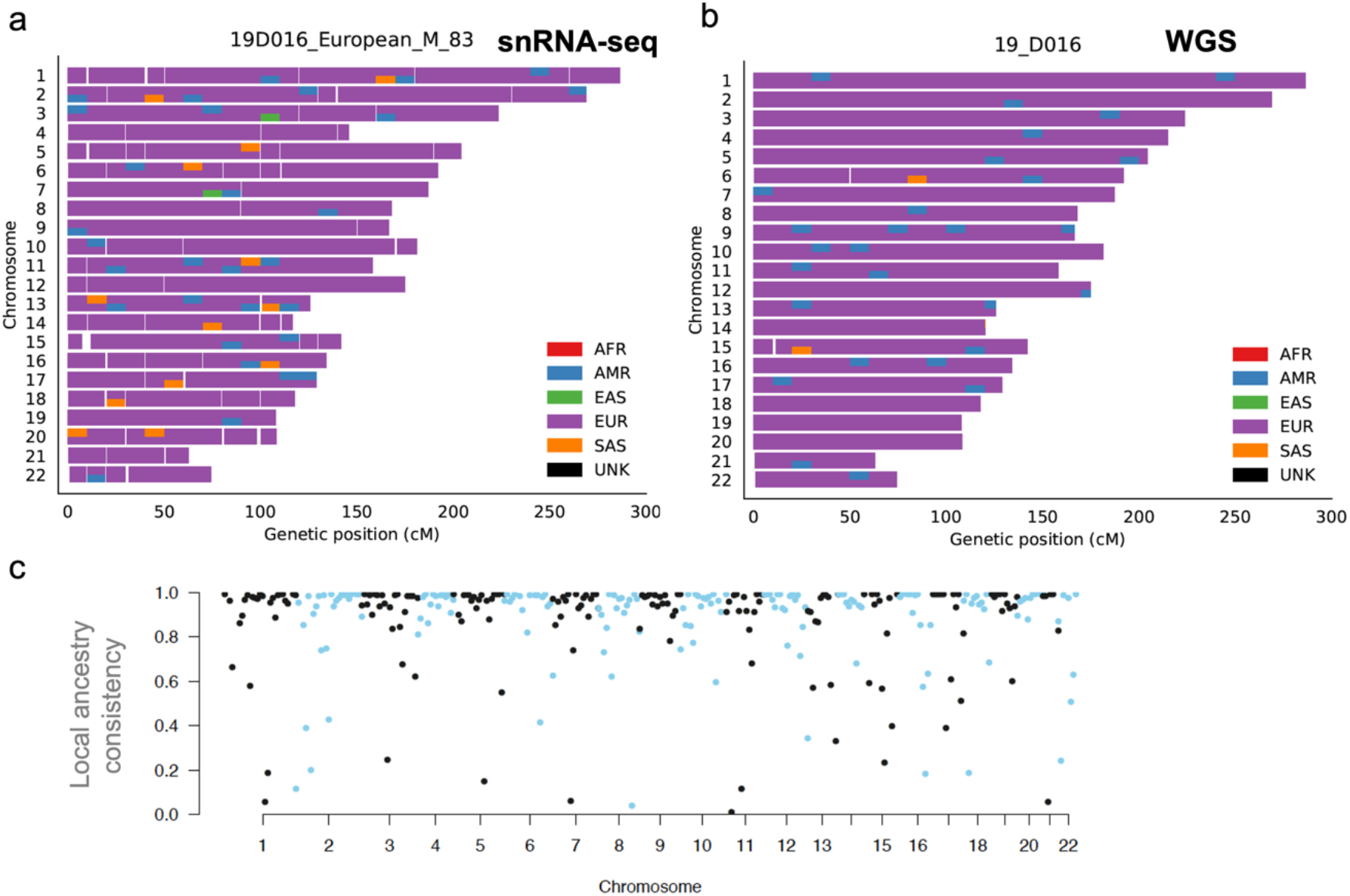
Local ancestry inference on 19D016 from the retina study. **a-b.** Local ancestry inference of a European sample 19D016 using genotypes from the snRNA-seq (**a**) and the WGS (**b**) data, respectively. The phased genotypes from 1KG3 are used as the reference for local ancestry inference. Colors in each chromosome denotes the inferred source ancestry with a bin size of 1cM.

**Fig.S4.**
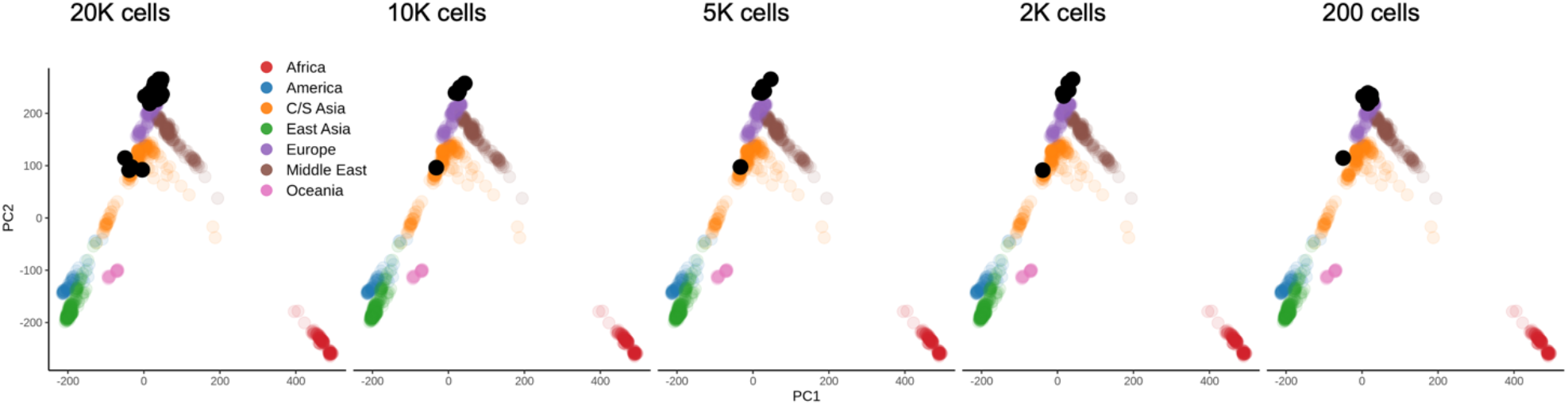
Genetic ancestry of the 4 retina samples using Monopogen genotypes derived from downsampled snRNA-seq data. The down-sampling scheme is the same as the ones in **Fig. 2e-f**. Colored dots represent the 897 reference individuals in the HGDP and black dots the retina samples.

**Fig.S5.**
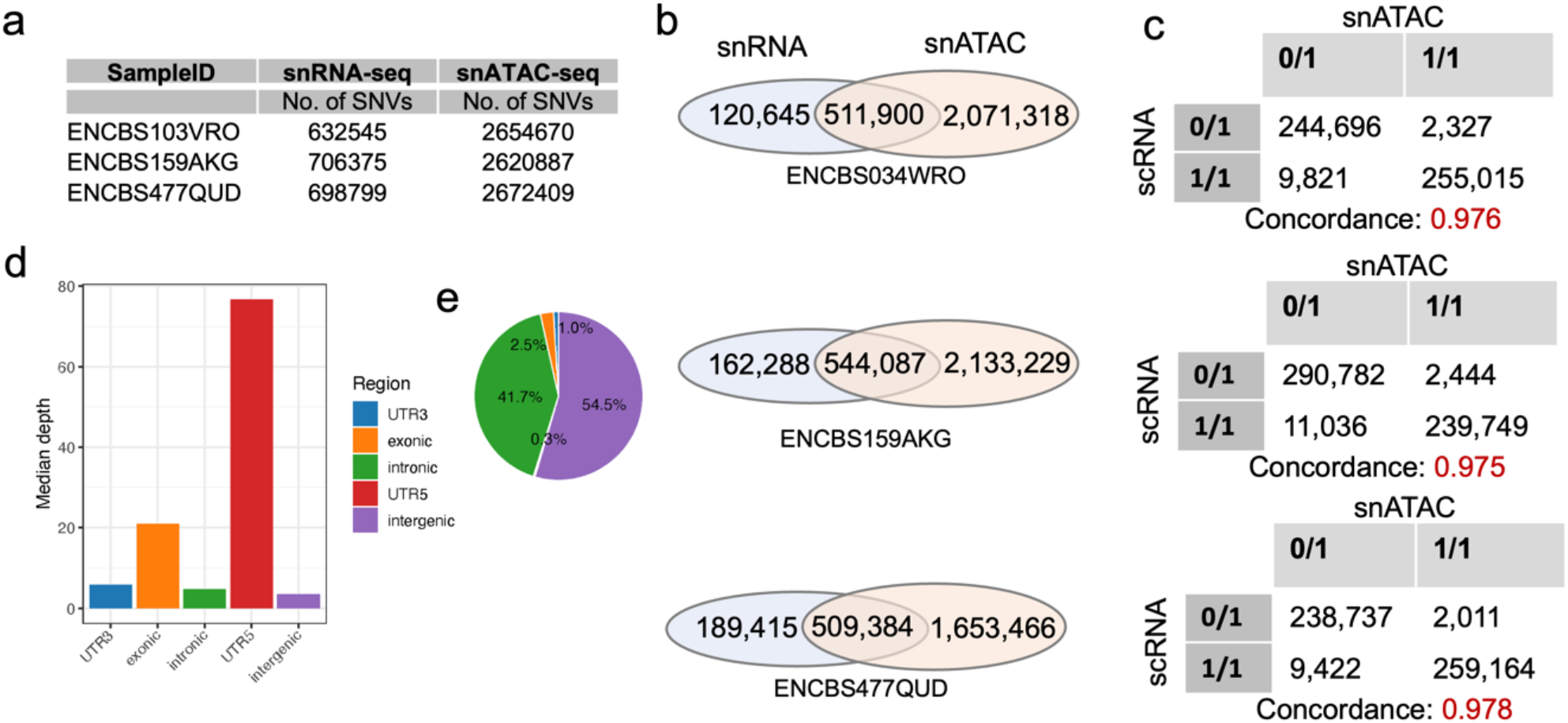
Genotyping concordance between the snRNA-seq and the snATAC-seq data from the same sample. **a,** The number of germline variants from randomly selected 3 heart left ventricle donors in the ENCODE project. Each sample has paired snRNA-seq and snATAC-seq measured. **b,** Variants overlapping between two modalities. **c.** Genotyping concordance between two modalities based on overlapping loci in **b**. **d-e,** Distribution of germline variants from Monopogen with median depth (**d**) and the percentage of variants (**e**) in each gene annotation category.

**Fig.S6.**
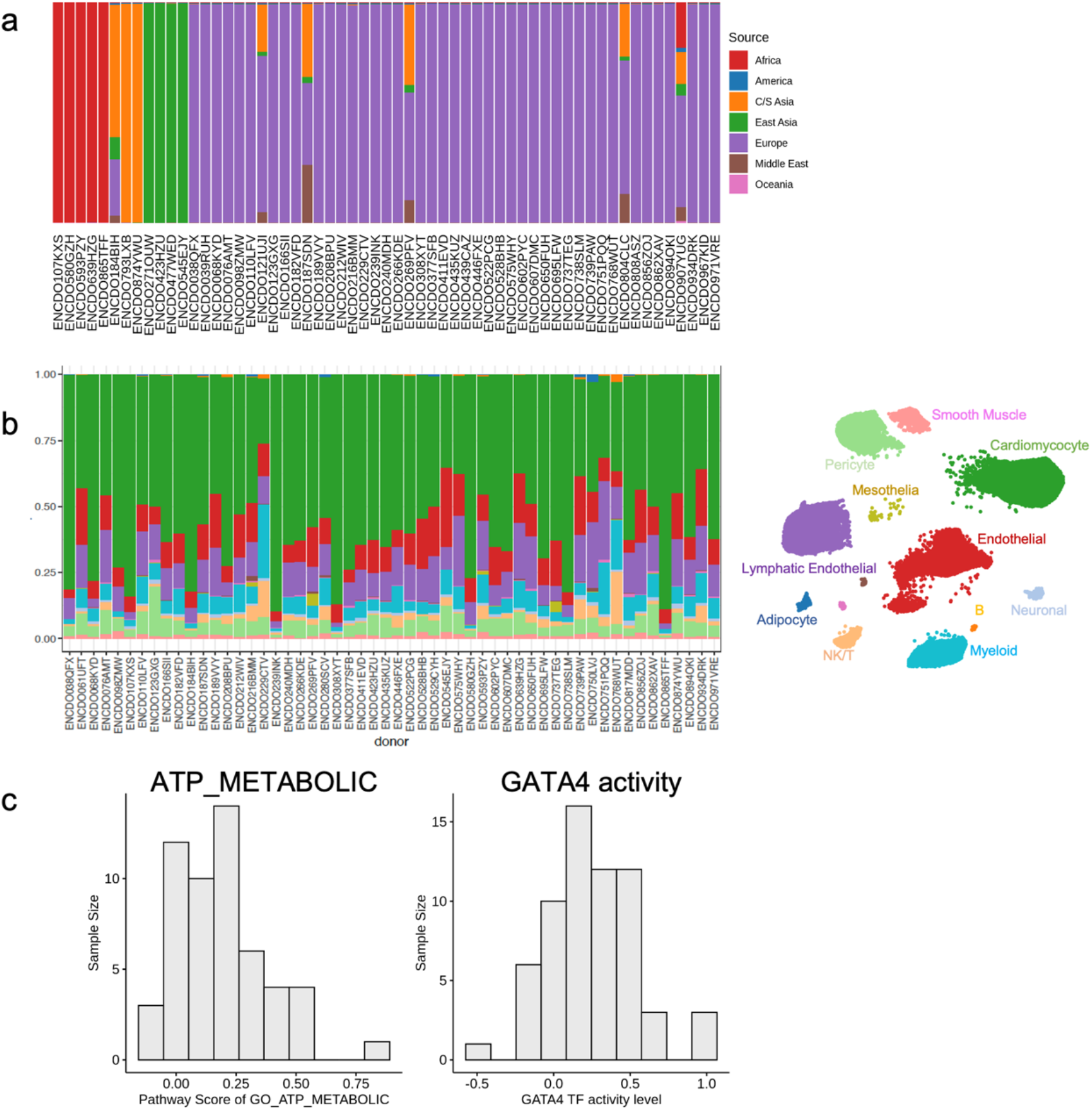
Sample information on 65 heart left ventricle donors in the ENCODE project. **a,** Inferred cell type composition by mapping cells to Azimuth heart database. **b,** Inferred ancestry components based on genotypes from Monopogen using ADMIXTURE. **c.** Histogram distribution of calculated pathway score GO_ATP_METABOLIC. **d.** Histogram distribution of calculated *GATA4* TF activity level from chromatin accessibility level.

**Fig.S7.**
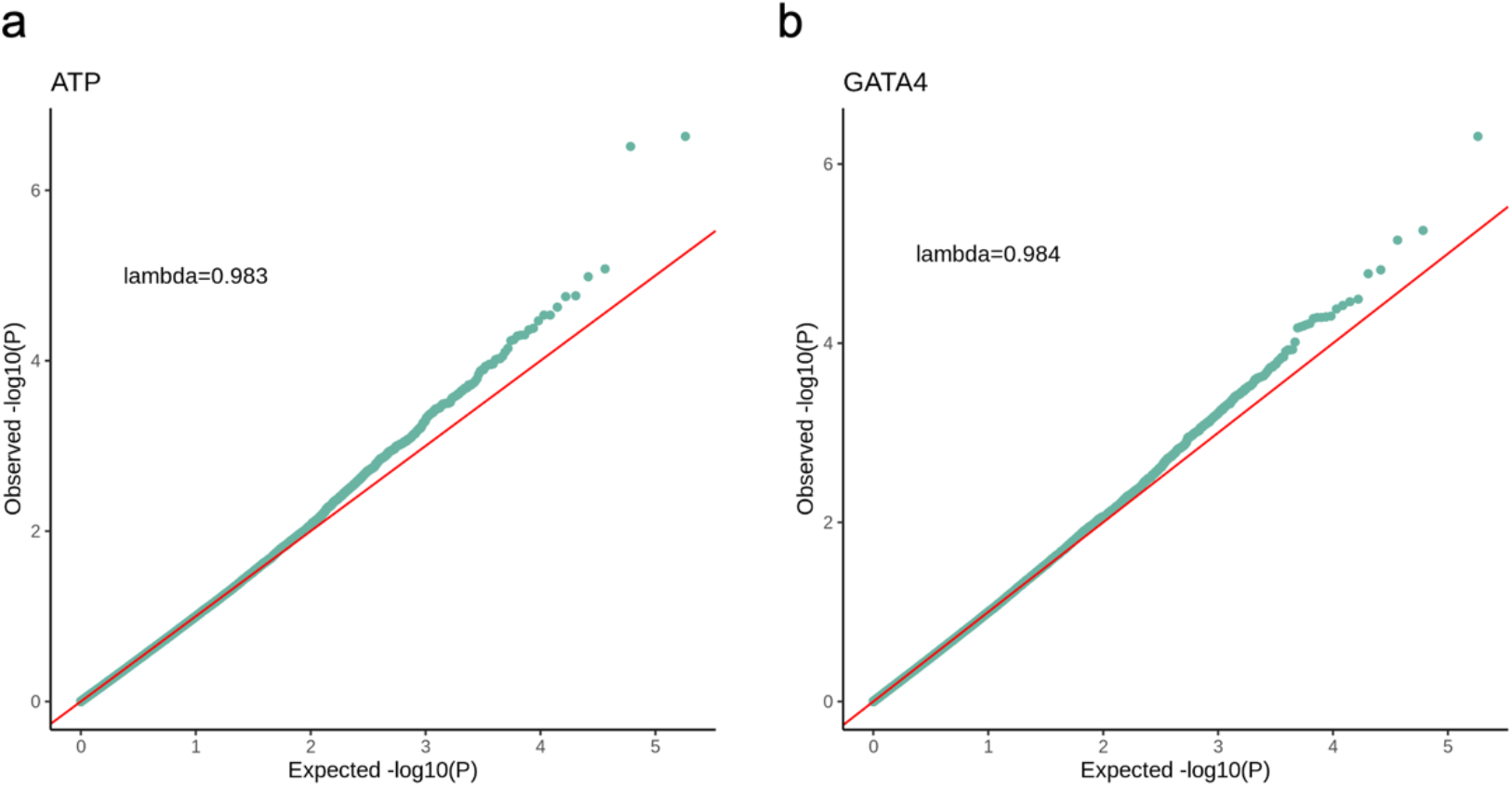
Quantile-Quantile plot for the association study of GO_ATP_METABOLIC (a) and GATA4 TF activity level (b). The inflation factor value is labeled on left top corner.

## Supplementary Tables

**Table S1.** Summary of variant calling on two benchmarking datasets for Monopogen.

**Table S2.** Summary of variant calling on two benchmarking datasets for Samtools and GATK.

**Table S3.** Summary of novel SNVs detected from Monopogen in 4 retina snRNA-seq samples (related to **Fig. S1b**).

**Table S4.** Summary of variant calling for HBCA, AIDA, and GTEx cohorts.

**Table S5.** Summary of variant calling from heart left ventricle samples. N.A: not available.

**Table S6.** Annotation of variants associated with GO_ATP_METABOLIC level with p-val<10^-5^.

**Table S7.** Annotation of variants associated with *GATA4* activity level with p-val<10^-5^.

## Notes

### Competing Interest Statement

The authors have declared no competing interest.

