## Supplementary Tables for "Monopogen: single nucleotide variant calling from single cell sequencing"

| | Study | SampleID | No.of cells | No. of reads | No. of reads per cell | Read length | No. of SNVs in WGS (W) | No. of SNVs in single cell sequencing (N | No. of SNVs overlapping (TP) | 0/1->1/1 | 1/1->0/1 | Recall (TP/W) | Precision (TP/N) | Genotyping accuracy | Overall accuracy |
|---|---|---|---|---|---|---|---|---|---|---|---|---|---|---|---|
| snRNA-seq | Retina tissue | 19D013 | 21016[a] | 722847191 | 33133 | 150bp | 4043921 | 830799 | 823045 | 2479 | 15114 | 0.203 | 0.991 | 0.979 | 0.970 |
| | | 19D014 | 20127 | 717949095 | 35670 | 150bp | 4069978 | 827731 | 820017 | 2476 | 15585 | 0.201 | 0.991 | 0.979 | 0.970 |
| | | 19D015 | 26395 | 619591103 | 23473 | 150bp | 4147482 | 885969 | 878586 | 2746 | 13108 | 0.212 | 0.992 | 0.982 | 0.974 |
| | | 19D016 | 25340 | 594626499 | 25340 | 150bp | 3925722 | 905395 | 892993 | 3276 | 12540 | 0.227 | 0.986 | 0.983 | 0.970 |
| sciATAC-seq | Transverse colon | ENCDO845WKR | 12122[b] | 67690360 | 5584 | 50bp | 3900140 | 1133545 | 1126660 | 4068 | 16289 | 0.289 | 0.994 | 0.982 | 0.976 |
| | | ENCDO793IXB | 12611 | 61044400 | 4840 | 50bp | 3728381 | 755565 | 752150 | 4038 | 28779 | 0.202 | 0.995 | 0.957 | 0.952 |

**Table S1.** Summary of SNV calling for two benchmarking datasets for Monopogen. [a]: cells with at least 500 genes expressed. [b]: cells with at least 500 peaks detected.

| | Platform | Study | SampleID | No. of SNVs in WGS (W) | No. of SNVs in single cells (N) | No. of SNVs overlapping (TP) | 0/1->1/1 | 1/1->0/1 | Recall (TP/W) | Precision (TP/N) | Genotyping accuracy | Overall accuracy |
|---|---|---|---|---|---|---|---|---|---|---|---|---|
| Samtools | snRNA-seq | Retina tissue | 19D013 | 4043921 | 1061214 | 768328 | 90076 | 3027 | 0.190 | 0.724 | 0.879 | 0.636 |
| | | | 19D014 | 4069978 | 1120514 | 828363 | 96285 | 3061 | 0.204 | 0.739 | 0.880 | 0.651 |
| | | | 19D015 | 4147482 | 1055049 | 840055 | 83022 | 2926 | 0.203 | 0.796 | 0.898 | 0.715 |
| | | | 19D016 | 3925722 | 1075721 | 826081 | 81474 | 3666 | 0.210 | 0.768 | 0.897 | 0.689 |
| | snATAC-seq | Transverse colon | ENCDO 845WKR | 3900140 | 188322 | 185654 | 64750 | 270 | 0.048 | 0.986 | 0.650 | 0.641 |
| | | | ENCDO 793IXB | 3728381 | 331891 | 329371 | 93192 | 40 | 0.088 | 0.992 | 0.717 | 0.711 |
| GATK | snRNA-seq | Retina tissue | 19D013 | 4043921 | 1068479 | 423234 | 3177 | 2724 | 0.105 | 0.396 | 0.986 | 0.391 |
| | | | 19D014 | 4069978 | 1114374 | 463466 | 3888 | 3016 | 0.114 | 0.416 | 0.985 | 0.410 |
| | | | 19D015 | 4147482 | 891334 | 455319 | 2395 | 2583 | 0.110 | 0.511 | 0.989 | 0.505 |
| | | | 19D016 | 3925722 | 926002 | 444972 | 2216 | 3238 | 0.113 | 0.481 | 0.988 | 0.475 |
| | snATAC-seq | Transverse colon | ENCDO 845WKR | 3900140 | 507967 | 464775 | 95537 | 627 | 0.119 | 0.915 | 0.793 | 0.726 |
| | | | ENCDO 793IXB | 3728381 | 292631 | 256275 | 53060 | 98 | 0.069 | 0.876 | 0.793 | 0.694 |

**Table S2.** Summary of SNV calling for two benchmarking datasets for Samtools and GATK.

| Fraction of clusters with SNV | SampleID | No. of SNVs called (N) | No. of SNVs overlapping in WGS (TP) | 0/1->0/1 | 0/1->1/1 | 1/1->0/1 | 1/1->1/1 | Precision (TP/N) | Genotyping accuracy | Overall accuracy |
|---|---|---|---|---|---|---|---|---|---|---|
| >0 | 19D013 | 129942 | 37674 | 37545 | 45 | 79 | 5 | 0.29 | 0.997 | 0.289 |
| >0.1 | 19D013 | 124875 | 37616 | 37497 | 36 | 79 | 4 | 0.301 | 0.997 | 0.300 |
| >0.2 | 19D013 | 106891 | 37228 | 37138 | 15 | 74 | 1 | 0.348 | 0.998 | 0.347 |
| >0.3 | 19D013 | 83565 | 36334 | 36261 | 4 | 69 | 0 | 0.435 | 0.998 | 0.434 |
| >0.4 | 19D013 | 64935 | 34919 | 34855 | 3 | 61 | 0 | 0.538 | 0.998 | 0.537 |
| >0.5 | 19D013 | 51398 | 32558 | 32499 | 2 | 57 | 0 | 0.633 | 0.998 | 0.632 |
| >0.6 | 19D013 | 41769 | 29500 | 29454 | 1 | 45 | 0 | 0.706 | 0.998 | 0.705 |
| >0.7 | 19D013 | 33213 | 25027 | 24985 | 1 | 41 | 0 | 0.754 | 0.998 | 0.752 |
| >0.8 | 19D013 | 24199 | 19079 | 19051 | 0 | 28 | 0 | 0.788 | 0.999 | 0.787 |
| >0.9 | 19D013 | 12247 | 10031 | 10020 | 0 | 11 | 0 | 0.819 | 0.999 | 0.818 |
| >0 | 19D014 | 118783 | 36182 | 35979 | 72 | 126 | 5 | 0.305 | 0.995 | 0.303 |
| >0.1 | 19D014 | 114956 | 36134 | 35941 | 62 | 126 | 5 | 0.314 | 0.995 | 0.312 |
| >0.2 | 19D014 | 99169 | 35821 | 35670 | 27 | 123 | 1 | 0.361 | 0.996 | 0.360 |
| >0.3 | 19D014 | 77232 | 35067 | 34950 | 3 | 114 | 0 | 0.454 | 0.997 | 0.453 |
| >0.4 | 19D014 | 59934 | 33806 | 33702 | 1 | 103 | 0 | 0.564 | 0.997 | 0.562 |
| >0.5 | 19D014 | 47528 | 31686 | 31593 | 0 | 93 | 0 | 0.667 | 0.997 | 0.665 |
| >0.6 | 19D014 | 38688 | 28707 | 28630 | 0 | 77 | 0 | 0.742 | 0.997 | 0.740 |
| >0.7 | 19D014 | 30583 | 24332 | 24265 | 0 | 67 | 0 | 0.796 | 0.997 | 0.794 |
| >0.8 | 19D014 | 22171 | 18406 | 18359 | 0 | 47 | 0 | 0.83 | 0.997 | 0.828 |
| >0.9 | 19D014 | 11150 | 9463 | 9432 | 0 | 31 | 0 | 0.849 | 0.997 | 0.846 |
| >0 | 19D015 | 69548 | 29467 | 29370 | 34 | 59 | 4 | 0.424 | 0.997 | 0.423 |
| >0.1 | 19D015 | 67710 | 29441 | 29351 | 27 | 59 | 4 | 0.435 | 0.997 | 0.434 |
| >0.2 | 19D015 | 59270 | 29201 | 29136 | 8 | 55 | 2 | 0.493 | 0.998 | 0.492 |
| >0.3 | 19D015 | 48496 | 28717 | 28663 | 1 | 52 | 1 | 0.592 | 0.998 | 0.591 |
| >0.4 | 19D015 | 40188 | 27902 | 27851 | 1 | 50 | 0 | 0.694 | 0.998 | 0.693 |
| >0.5 | 19D015 | 34524 | 26581 | 26537 | 0 | 44 | 0 | 0.77 | 0.998 | 0.768 |
| >0.6 | 19D015 | 30417 | 24877 | 24836 | 0 | 41 | 0 | 0.818 | 0.998 | 0.816 |
| >0.7 | 19D015 | 26685 | 22655 | 22622 | 0 | 33 | 0 | 0.849 | 0.999 | 0.848 |
| >0.8 | 19D015 | 20575 | 18207 | 18181 | 0 | 26 | 0 | 0.885 | 0.999 | 0.884 |
| >0.9 | 19D015 | 12066 | 10952 | 10939 | 0 | 13 | 0 | 0.908 | 0.999 | 0.907 |
| >0 | 19D016 | 79931 | 30432 | 30258 | 47 | 123 | 4 | 0.381 | 0.994 | 0.379 |
| >0.1 | 19D016 | 77746 | 30394 | 30230 | 37 | 123 | 4 | 0.391 | 0.995 | 0.389 |
| >0.2 | 19D016 | 67693 | 30187 | 30050 | 13 | 120 | 4 | 0.446 | 0.996 | 0.444 |
| >0.3 | 19D016 | 54813 | 29641 | 29520 | 3 | 117 | 1 | 0.541 | 0.996 | 0.539 |
| >0.4 | 19D016 | 44836 | 28825 | 28712 | 0 | 112 | 1 | 0.643 | 0.996 | 0.640 |
| >0.5 | 19D016 | 38119 | 27556 | 27449 | 0 | 106 | 1 | 0.723 | 0.996 | 0.720 |
| >0.6 | 19D016 | 33143 | 25756 | 25663 | 0 | 92 | 1 | 0.777 | 0.996 | 0.774 |
| >0.7 | 19D016 | 28721 | 23278 | 23196 | 0 | 82 | 0 | 0.81 | 0.996 | 0.807 |
| >0.8 | 19D016 | 22224 | 18922 | 18859 | 0 | 63 | 0 | 0.851 | 0.997 | 0.848 |
| >0.9 | 19D016 | 12548 | 10972 | 10942 | 0 | 30 | 0 | 0.874 | 0.997 | 0.871 |

**Table S3.** Summary of novel SNVs detected from Monopogen in 4 retina snRNA-seq samples (related to **Fig. S1b**).

| | SampleID | No. of cells | No. of reads | No. of reads per cell | Read length | No. of germline SNVs |
|---|---|---|---|---|---|---|
| Asia Immune Diversity Study | JP_H045 | 1569 | 59239054 | 37755 | 150bp | 121313 |
| | JP_H046 | 1189 | 39486450 | 33209 | 150bp | 99964 |
| | JP_H047 | 1221 | 41068349 | 33635 | 150bp | 110858 |
| | JP_H048 | 1142 | 37410391 | 32758 | 150bp | 93531 |
| | JP_H137 | 1197 | 45362685 | 37896 | 150bp | 124220 |
| | JP_H146 | 1117 | 44239330 | 39605 | 150bp | 112796 |
| | JP_H148 | 1112 | 44318390 | 39854 | 150bp | 120655 |
| | JP_H149 | 1411 | 57020159 | 40411 | 150bp | 132721 |
| | KR_H001 | 1264 | 57476150 | 45471 | 150bp | 163256 |
| | KR_H002 | 840 | 35435379 | 42184 | 150bp | 118850 |
| | KR_H004 | 821 | 34368945 | 41862 | 150bp | 128936 |
| | KR_H005 | 918 | 35884600 | 39089 | 150bp | 114778 |
| | KR_H160 | 1279 | 47380301 | 37044 | 150bp | 120229 |
| | KR_H161 | 1573 | 57235627 | 36386 | 150bp | 131808 |
| | KR_H164 | 1573 | 57235627 | 36386 | 150bp | 120423 |
| | KR_H165 | 1602 | 52719939 | 32908 | 150bp | 103942 |
| | Lonza_3038016 | 1300 | 40815193 | 31396 | 150bp | 102320 |
| | Lonza_3038097 | 999 | 44136732 | 44180 | 150bp | 98936 |
| | Lonza_3038099 | 1049 | 48096207 | 45849 | 150bp | 77076 |
| | Lonza_3038306 | 761 | 23257052 | 30561 | 150bp | 82056 |
| Human Breast Cell Atlas | HBCA02C | 1709 | 324492736 | 189872 | 125bp | 256088 |
| | HBCA02i | 2232 | 331161319 | 148369 | 125bp | 369781 |
| | HBCA03C | 1865 | 356381426 | 191089 | 125bp | 471241 |
| | HBCA03Ccryo | 1525 | 348639606 | 228616 | 125bp | 310490 |
| | HBCA03i | 1470 | 356132757 | 242267 | 125bp | 442943 |
| | HBCA04C | 1810 | 357352385 | 197432 | 125bp | 380657 |
| | HBCA04i | 759 | 349784479 | 460849 | 125bp | 390222 |
| | HBCA04T | 1608 | 348689072 | 216846 | 125bp | 352607 |
| | HBCA05C | 464 | 362981567 | 782287 | 125bp | 441287 |
| | HBCA05Ccryo | 1145 | 349034669 | 304833 | 125bp | 293419 |
| | HBCA05i | 1663 | 348892182 | 209796 | 125bp | 430660 |
| | HBCA06C | 1045 | 330866587 | 316618 | 125bp | 390307 |
| | HBCA07C | 1405 | 320441257 | 228072 | 125bp | 379910 |
| | HBCA07i | 2451 | 319876418 | 130508 | 125bp | 429340 |
| | HBCA07T | 2226 | 314283146 | 141187 | 125bp | 454028 |
| | HBCA09C | 1231 | 319872861 | 259847 | 125bp | 385241 |
| | HBCA09i | 2027 | 323445690 | 159568 | 125bp | 314550 |
| | HBCA10C | 1466 | 362133748 | 247021 | 125bp | 267024 |
| | HBCA10Ccryo | 1297 | 348249946 | 268504 | 125bp | 300322 |
| | HBCA10i | 891 | 345961110 | 388284 | 125bp | 237928 |
| GETx | GTEX-13N11 | 5226 | 119605363 | 22887 | 100bp | 300538 |
| | GTEX-144GM | 5289 | 89603310 | 16941 | 100bp | 181382 |
| | GTEX-15CHR | 3508 | 115170743 | 32830 | 100bp | 155046 |
| | GTEX-15RIE | 5001 | 114957894 | 22987 | 100bp | 205432 |
| | GTEX-15SB6 | 4819 | 128624204 | 26691 | 100bp | 304507 |
| | GTEX-16BQI | 5269 | 116701973 | 22148 | 100bp | 211307 |
| | GTEX-1ICG6 | 4148 | 109050028 | 26289 | 100bp | 216682 |

**Table S4.** Summary of SNV calling for AIDA, HBCA, and GTEx cohorts.

| SampleID | No. of SNVs in snRNA-seq | No. of SNVs in snATAC-seq | SampleID | No. of SNVs in snRNA-seq | No. of SNVs in snATAC-seq |
|---|---|---|---|---|---|
| ENCDO738SLM | 537076 | 2602120 | ENCDO271OUW | N.A | 2468503 |
| ENCDO575WHY | 788194 | 2599335 | ENCDO793LXB | N.A | 2590288 |
| ENCDO377SFB | 646028 | 2595870 | ENCDO189VVY | 698813 | 2601870 |
| ENCDO411EVD | 790506 | 2592981 | ENCDO123GXG | 726431 | 2594985 |
| ENCDO650FUH | 949924 | 2609254 | ENCDO266KDE | 1031286 | 2591638 |
| ENCDO229CTV | 998609 | 2591653 | ENCDO182VFD | 942648 | 2601761 |
| ENCDO934DRK | 984814 | 2603980 | ENCDO308XYT | 704328 | 2600948 |
| ENCDO269PFV | 1176624 | 2593567 | ENCDO239INK | 802663 | 2591464 |
| ENCDO107KXS | 954463 | 2460631 | ENCDO971VRE | 939773 | 2587254 |
| ENCDO435KUZ | 903239 | 2595532 | ENCDO423HZU | 892430 | 2459095 |
| ENCDO184BIH | 760618 | 2530747 | ENCDO593PZY | 1061712 | 2453326 |
| ENCDO602PYC | 752144 | 2594180 | ENCDO862XAV | 719859 | 2610909 |
| ENCDO038QFX | 676060 | 2596614 | ENCDO212WIV | 642434 | 2579584 |
| ENCDO216BMM | 707303 | 2619222 | ENCDO280SCV | 671894 | 2599335 |
| ENCDO528BHB | 780800 | 2596634 | ENCDO817MDD | 693944 | 2623213 |
| ENCDO737TEG | 881202 | 2608524 | ENCDO607DMC | 685762 | 2587715 |
| ENCDO061UFT | 759082 | 2617048 | ENCDO446FXE | 690928 | 2597200 |
| ENCDO751PQQ | 1063673 | 2608314 | ENCDO110LFV | 710207 | 2599868 |
| ENCDO874YWU | 632607 | 2587876 | ENCDO477WED | N.A | 2481147 |
| ENCDO639HZG | 706263 | 2462721 | ENCDO768WUT | 680723 | 2599976 |
| ENCDO545EJY | 804425 | 2446069 | ENCDO208BPU | 649945 | 2589330 |
| ENCDO739PAW | 592706 | 2602392 | ENCDO098ZMW | 634326 | 2589542 |
| ENCDO750LVJ | 1038750 | 2589178 | ENCDO695LFW | 702332 | 2618752 |
| ENCDO580GZH | 1085464 | 2466424 | ENCDO907YUG | N.A | 2964332 |
| ENCDO894OKI | 881239 | 2588923 | ENCDO967KID | N.A | 2596910 |
| ENCDO187SDN | 908590 | 2603997 | ENCDO804CLC | N.A | 2585524 |
| ENCDO865TFF | 790664 | 2459535 | ENCDO926KEV | N.A | 2693594 |
| ENCDO076AMT | 418685 | 2601072 | ENCDO808ASZ | N.A | 2627648 |
| ENCDO068KYD | 886965 | 2599461 | ENCDO439CAZ | N.A | 2605541 |
| ENCDO522PCG | 706117 | 2613923 | ENCDO039RUH | N.A | 2607601 |
| ENCDO529CYH | 669832 | 2615742 | ENCDO121UJI | N.A | 2633926 |
| ENCDO856ZOJ | 888770 | 2600768 | ENCDO240MDH | 639683 | 2597854 |
| | | | ENCDO166SII | 658079 | 2587348 |

**Table S5.** Summary of SNV calling from heart left ventricle samples. N.A: not available.

| rsID | CHR | POS | A1 | A2 | AF1 | BETA | SE | P | distancetoFeature | symbol |
|---|---|---|---|---|---|---|---|---|---|---|
| rs12748496 | 1 | 59020879 | A | G | 0.265 | 0.168 | 0.038 | 8.08E-06 | 87235 | LINC01358 |
| rs12124171 | 1 | 80188832 | G | A | 0.204 | -0.206 | 0.042 | 8.70E-07 | -346924 | LINC01781 |
| rs10159130 | 1 | 80192436 | C | T | 0.204 | -0.206 | 0.042 | 8.70E-07 | -343320 | LINC01781 |
| rs11162948 | 1 | 80192975 | C | T | 0.204 | -0.206 | 0.042 | 8.70E-07 | -342781 | LINC01781 |
| rs17104484 | 1 | 80193168 | C | A | 0.204 | -0.206 | 0.042 | 8.70E-07 | -342588 | LINC01781 |
| rs11162949 | 1 | 80195006 | G | A | 0.204 | -0.206 | 0.042 | 8.70E-07 | -340750 | LINC01781 |
| rs10782722 | 1 | 80197213 | G | A | 0.204 | -0.206 | 0.042 | 8.70E-07 | -338543 | LINC01781 |
| rs10782723 | 1 | 80197498 | C | G | 0.204 | -0.206 | 0.042 | 8.70E-07 | -338258 | LINC01781 |
| rs1529999 | 2 | 118476740 | T | C | 0.500 | -0.128 | 0.028 | 5.87E-06 | 370939 | EN1 |
| rs4849701 | 2 | 118478920 | C | T | 0.510 | -0.139 | 0.028 | 6.15E-07 | 368759 | EN1 |
| rs6542466 | 2 | 118481948 | A | G | 0.520 | -0.133 | 0.028 | 1.72E-06 | 365731 | EN1 |
| rs4848510 | 2 | 118482581 | A | C | 0.449 | -0.129 | 0.028 | 4.12E-06 | 365098 | EN1 |
| rs1344896 | 2 | 118492854 | T | C | 0.449 | -0.129 | 0.028 | 4.12E-06 | 354825 | EN1 |
| rs6752749 | 2 | 118499583 | G | C | 0.449 | -0.129 | 0.028 | 4.12E-06 | 348096 | EN1 |
| rs13009652 | 2 | 138959515 | C | T | 0.265 | 0.179 | 0.040 | 7.79E-06 | -179166 | NXPH2 |
| rs1710827 | 2 | 235760076 | A | G | 0.296 | 0.160 | 0.030 | 6.52E-08 | 254324 | AGAP1-IT1 |
| rs2696393 | 2 | 235760409 | G | A | 0.296 | 0.160 | 0.030 | 6.52E-08 | 254657 | AGAP1-IT1 |
| rs2696394 | 2 | 235760459 | C | G | 0.296 | 0.160 | 0.030 | 6.52E-08 | 254707 | AGAP1-IT1 |
| rs2696395 | 2 | 235760542 | A | G | 0.296 | 0.160 | 0.030 | 6.52E-08 | 254790 | AGAP1-IT1 |
| rs2696396 | 2 | 235760568 | G | A | 0.296 | 0.160 | 0.030 | 6.52E-08 | 254816 | AGAP1-IT1 |
| rs2675142 | 2 | 235760894 | G | A | 0.296 | 0.160 | 0.030 | 6.52E-08 | 255142 | AGAP1-IT1 |
| rs2696397 | 2 | 235761153 | T | C | 0.296 | 0.160 | 0.030 | 6.52E-08 | 255401 | AGAP1-IT1 |
| rs72975265 | 2 | 235761194 | G | A | 0.296 | 0.160 | 0.030 | 6.52E-08 | 255442 | AGAP1-IT1 |
| rs12469618 | 2 | 235761287 | G | A | 0.296 | 0.160 | 0.030 | 6.52E-08 | 255535 | AGAP1-IT1 |
| rs12469659 | 2 | 235761387 | G | A | 0.296 | 0.160 | 0.030 | 6.52E-08 | 255635 | AGAP1-IT1 |
| rs6714660 | 2 | 235761752 | C | T | 0.296 | 0.160 | 0.030 | 6.52E-08 | 256000 | AGAP1-IT1 |
| rs6715112 | 2 | 235762292 | G | A | 0.296 | 0.160 | 0.030 | 6.52E-08 | 256540 | AGAP1-IT1 |
| rs6718263 | 2 | 235762347 | G | T | 0.296 | 0.160 | 0.030 | 6.52E-08 | 256595 | AGAP1-IT1 |
| rs2696403 | 2 | 235762546 | A | G | 0.296 | 0.160 | 0.030 | 6.52E-08 | 256794 | AGAP1-IT1 |
| rs2696402 | 2 | 235762632 | T | C | 0.296 | 0.160 | 0.030 | 6.52E-08 | 256880 | AGAP1-IT1 |
| rs2696401 | 2 | 235762650 | T | C | 0.286 | 0.166 | 0.031 | 8.09E-08 | 256898 | AGAP1-IT1 |
| rs2675141 | 2 | 235762859 | C | T | 0.296 | 0.160 | 0.030 | 6.52E-08 | 257107 | AGAP1-IT1 |
| rs3896943 | 2 | 235762882 | A | G | 0.296 | 0.160 | 0.030 | 6.52E-08 | 257130 | AGAP1-IT1 |
| rs938492 | 2 | 235762995 | C | G | 0.306 | 0.157 | 0.030 | 2.33E-07 | 257243 | AGAP1-IT1 |
| rs938491 | 2 | 235763052 | G | A | 0.316 | 0.153 | 0.031 | 7.22E-07 | 257300 | AGAP1-IT1 |
| rs2696405 | 2 | 235763462 | C | T | 0.306 | 0.157 | 0.030 | 2.33E-07 | 257710 | AGAP1-IT1 |
| rs2696406 | 2 | 235763593 | G | T | 0.306 | 0.157 | 0.030 | 2.33E-07 | 257841 | AGAP1-IT1 |
| rs2696407 | 2 | 235763618 | C | T | 0.306 | 0.157 | 0.030 | 2.33E-07 | 257866 | AGAP1-IT1 |
| rs552845949 | 2 | 235764030 | G | T | 0.286 | 0.157 | 0.032 | 8.88E-07 | 258278 | AGAP1-IT1 |
| rs2675139 | 2 | 235764071 | G | A | 0.306 | 0.157 | 0.030 | 2.33E-07 | 258319 | AGAP1-IT1 |
| rs2696409 | 2 | 235764087 | C | T | 0.316 | 0.156 | 0.031 | 3.03E-07 | 258335 | AGAP1-IT1 |
| rs2696410 | 2 | 235764169 | A | G | 0.306 | 0.157 | 0.030 | 2.33E-07 | 258417 | AGAP1-IT1 |
| rs2675138 | 2 | 235764317 | T | C | 0.306 | 0.157 | 0.030 | 2.33E-07 | 258565 | AGAP1-IT1 |
| rs2675136 | 2 | 235764447 | T | C | 0.306 | 0.157 | 0.030 | 2.33E-07 | 258695 | AGAP1-IT1 |
| rs2675135 | 2 | 235764536 | G | C | 0.296 | 0.161 | 0.032 | 3.06E-07 | 258784 | AGAP1-IT1 |
| rs6712556 | 2 | 235765406 | T | C | 0.306 | 0.157 | 0.030 | 2.33E-07 | 259654 | AGAP1-IT1 |
| rs6712558 | 2 | 235765411 | T | C | 0.306 | 0.157 | 0.030 | 2.33E-07 | 259659 | AGAP1-IT1 |
| rs2675134 | 2 | 235765760 | C | T | 0.306 | 0.157 | 0.030 | 2.33E-07 | 260008 | AGAP1-IT1 |
| rs2675133 | 2 | 235766294 | T | C | 0.296 | 0.161 | 0.032 | 3.06E-07 | 260542 | AGAP1-IT1 |
| rs2696386 | 2 | 235766387 | A | G | 0.296 | 0.161 | 0.032 | 3.06E-07 | 260635 | AGAP1-IT1 |
| rs2696387 | 2 | 235766438 | T | G | 0.306 | 0.157 | 0.030 | 2.33E-07 | 260686 | AGAP1-IT1 |
| rs2675132 | 2 | 235767018 | G | A | 0.296 | 0.161 | 0.032 | 3.06E-07 | 261266 | AGAP1-IT1 |
| rs2675131 | 2 | 235767377 | T | C | 0.296 | 0.161 | 0.032 | 3.06E-07 | 261625 | AGAP1-IT1 |
| rs2696388 | 2 | 235767450 | C | T | 0.296 | 0.161 | 0.032 | 3.06E-07 | 261698 | AGAP1-IT1 |
| rs2675130 | 2 | 235767464 | C | T | 0.296 | 0.161 | 0.032 | 3.06E-07 | 261712 | AGAP1-IT1 |
| rs2696389 | 2 | 235767675 | G | C | 0.296 | 0.161 | 0.032 | 3.06E-07 | 261923 | AGAP1-IT1 |

| | | | | | | | | | | |
|---|---|---|---|---|---|---|---|---|---|---|
| rs2675129 | 2 | 235767934 | G | A | 0.296 | 0.161 | 0.032 | 3.06E-07 | 262182 | AGAP1-IT1 |
| rs2696391 | 2 | 235768052 | T | G | 0.296 | 0.161 | 0.032 | 3.06E-07 | 262300 | AGAP1-IT1 |
| rs2696392 | 2 | 235768704 | G | C | 0.296 | 0.161 | 0.032 | 3.06E-07 | 262952 | AGAP1-IT1 |
| rs2675128 | 2 | 235769187 | G | A | 0.296 | 0.161 | 0.032 | 3.06E-07 | 263435 | AGAP1-IT1 |
| rs1728290 | 2 | 235770056 | G | A | 0.296 | 0.161 | 0.032 | 3.06E-07 | 264304 | AGAP1-IT1 |
| rs55907328 | 2 | 235770216 | C | T | 0.296 | 0.161 | 0.032 | 3.06E-07 | 264464 | AGAP1-IT1 |
| rs55749012 | 2 | 235770415 | A | G | 0.296 | 0.161 | 0.032 | 3.06E-07 | 264663 | AGAP1-IT1 |
| rs55797253 | 2 | 235770425 | G | A | 0.296 | 0.161 | 0.032 | 3.06E-07 | 264673 | AGAP1-IT1 |
| rs3108498 | 2 | 235770557 | C | T | 0.306 | 0.157 | 0.030 | 2.33E-07 | 264805 | AGAP1-IT1 |
| rs1710823 | 2 | 235770800 | T | C | 0.296 | 0.161 | 0.032 | 3.06E-07 | 265048 | AGAP1-IT1 |
| rs1710822 | 2 | 235770942 | A | C | 0.306 | 0.157 | 0.030 | 2.33E-07 | 265190 | AGAP1-IT1 |
| rs2675127 | 2 | 235771440 | C | G | 0.296 | 0.161 | 0.032 | 3.06E-07 | 265688 | AGAP1-IT1 |
| rs2675126 | 2 | 235771482 | C | T | 0.296 | 0.161 | 0.032 | 3.06E-07 | 265730 | AGAP1-IT1 |
| rs2675125 | 2 | 235771605 | G | A | 0.296 | 0.161 | 0.032 | 3.06E-07 | 265853 | AGAP1-IT1 |
| rs55926594 | 2 | 235771838 | T | C | 0.296 | 0.161 | 0.032 | 3.06E-07 | 266086 | AGAP1-IT1 |
| rs56071605 | 2 | 235771872 | T | C | 0.296 | 0.161 | 0.032 | 3.06E-07 | 266120 | AGAP1-IT1 |
| rs117993231 | 2 | 235771999 | T | C | 0.286 | 0.162 | 0.033 | 1.36E-06 | 266247 | AGAP1-IT1 |
| rs115957313 | 2 | 235772047 | G | A | 0.296 | 0.161 | 0.032 | 3.06E-07 | 266295 | AGAP1-IT1 |
| rs3108499 | 2 | 235772298 | G | A | 0.296 | 0.161 | 0.032 | 3.06E-07 | 266546 | AGAP1-IT1 |
| rs3106791 | 2 | 235772312 | T | C | 0.296 | 0.161 | 0.032 | 3.06E-07 | 266560 | AGAP1-IT1 |
| rs3106792 | 2 | 235772645 | T | C | 0.296 | 0.161 | 0.032 | 3.06E-07 | 266893 | AGAP1-IT1 |
| rs3108501 | 2 | 235772820 | A | G | 0.296 | 0.161 | 0.032 | 3.06E-07 | 267068 | AGAP1-IT1 |
| rs72977127 | 2 | 235773027 | C | T | 0.296 | 0.161 | 0.032 | 3.06E-07 | 267275 | AGAP1-IT1 |
| rs2696413 | 2 | 235773216 | T | G | 0.296 | 0.161 | 0.032 | 3.06E-07 | 267464 | AGAP1-IT1 |
| rs2696412 | 2 | 235774212 | C | T | 0.296 | 0.161 | 0.032 | 3.06E-07 | 268460 | AGAP1-IT1 |
| rs2696411 | 2 | 235774270 | C | T | 0.296 | 0.161 | 0.032 | 3.06E-07 | 268518 | AGAP1-IT1 |
| rs2249999 | 2 | 235774743 | A | C | 0.296 | 0.161 | 0.032 | 3.06E-07 | 268991 | AGAP1-IT1 |
| rs1356223 | 2 | 235774940 | G | A | 0.296 | 0.161 | 0.032 | 3.06E-07 | 269188 | AGAP1-IT1 |
| rs1400147 | 2 | 235775059 | G | A | 0.296 | 0.161 | 0.032 | 3.06E-07 | 269307 | AGAP1-IT1 |
| rs2292704 | 2 | 235775848 | C | T | 0.296 | 0.161 | 0.032 | 3.06E-07 | 270096 | AGAP1-IT1 |
| rs2292705 | 2 | 235775902 | C | T | 0.296 | 0.161 | 0.032 | 3.06E-07 | 270150 | AGAP1-IT1 |
| rs12465436 | 2 | 235776372 | C | T | 0.306 | 0.161 | 0.032 | 4.11E-07 | 270620 | AGAP1-IT1 |
| rs12468185 | 2 | 235776373 | C | G | 0.306 | 0.161 | 0.032 | 4.11E-07 | 270621 | AGAP1-IT1 |
| rs12468221 | 2 | 235776470 | C | G | 0.296 | 0.161 | 0.032 | 3.06E-07 | 270718 | AGAP1-IT1 |
| rs72981174 | 2 | 235776777 | A | C | 0.296 | 0.161 | 0.032 | 3.06E-07 | 271025 | AGAP1-IT1 |
| rs72981176 | 2 | 235776888 | A | G | 0.296 | 0.161 | 0.032 | 3.06E-07 | 271136 | AGAP1-IT1 |
| rs113739109 | 2 | 235777087 | C | G | 0.296 | 0.161 | 0.032 | 3.06E-07 | 271335 | AGAP1-IT1 |
| rs138493201 | 2 | 235777118 | G | A | 0.306 | 0.161 | 0.032 | 4.11E-07 | 271366 | AGAP1-IT1 |
| rs12151454 | 2 | 235777185 | C | T | 0.296 | 0.161 | 0.032 | 3.06E-07 | 271433 | AGAP1-IT1 |
| rs72981179 | 2 | 235777749 | T | C | 0.296 | 0.161 | 0.032 | 3.06E-07 | 271997 | AGAP1-IT1 |
| rs55961214 | 2 | 235778520 | A | G | 0.265 | 0.168 | 0.033 | 2.45E-07 | 272768 | AGAP1-IT1 |
| rs61402262 | 2 | 235778549 | T | C | 0.296 | 0.161 | 0.032 | 3.06E-07 | 272797 | AGAP1-IT1 |
| rs12470717 | 2 | 235778877 | A | G | 0.296 | 0.161 | 0.032 | 3.06E-07 | 273125 | AGAP1-IT1 |
| rs12463796 | 2 | 235779111 | T | C | 0.296 | 0.161 | 0.032 | 3.06E-07 | 273359 | AGAP1-IT1 |
| rs13400202 | 2 | 235780146 | G | A | 0.296 | 0.161 | 0.032 | 3.06E-07 | 274394 | AGAP1-IT1 |
| rs12472838 | 2 | 235780401 | A | G | 0.296 | 0.161 | 0.032 | 3.06E-07 | 274649 | AGAP1-IT1 |
| rs12470048 | 2 | 235780439 | G | T | 0.296 | 0.161 | 0.032 | 3.06E-07 | 274687 | AGAP1-IT1 |
| rs60591163 | 2 | 235781589 | C | T | 0.296 | 0.161 | 0.032 | 3.06E-07 | 275837 | AGAP1-IT1 |
| rs72981189 | 2 | 235781867 | A | G | 0.296 | 0.161 | 0.032 | 3.06E-07 | 276115 | AGAP1-IT1 |
| rs963846 | 2 | 235783052 | G | C | 0.684 | -0.150 | 0.031 | 1.69E-06 | 277300 | AGAP1-IT1 |
| rs373964138 | 2 | 235783214 | A | T | 0.296 | 0.161 | 0.032 | 3.06E-07 | 277462 | AGAP1-IT1 |
| rs1878154 | 2 | 235783338 | G | A | 0.296 | 0.161 | 0.032 | 3.06E-07 | 277586 | AGAP1-IT1 |
| rs12466588 | 2 | 235783594 | G | A | 0.296 | 0.161 | 0.032 | 3.06E-07 | 277842 | AGAP1-IT1 |
| rs72981192 | 2 | 235783821 | T | C | 0.296 | 0.161 | 0.032 | 3.06E-07 | 278069 | AGAP1-IT1 |
| rs59832814 | 2 | 235785072 | G | A | 0.296 | 0.161 | 0.032 | 3.06E-07 | 279320 | AGAP1-IT1 |
| rs56009264 | 2 | 235785796 | G | T | 0.296 | 0.161 | 0.032 | 3.06E-07 | 280044 | AGAP1-IT1 |
| rs12469834 | 2 | 235786022 | G | A | 0.296 | 0.161 | 0.032 | 3.06E-07 | 280270 | AGAP1-IT1 |

| | | | | | | | | | | |
|---|---|---|---|---|---|---|---|---|---|---|
| rs57957542 | 2 | 235786605 | A | G | 0.296 | 0.161 | 0.032 | 3.06E-07 | 280853 | AGAP1-IT1 |
| rs61499079 | 2 | 235786797 | G | C | 0.296 | 0.161 | 0.032 | 3.06E-07 | 281045 | AGAP1-IT1 |
| rs1878155 | 2 | 235787192 | T | G | 0.296 | 0.161 | 0.032 | 3.06E-07 | 281440 | AGAP1-IT1 |
| rs12472982 | 2 | 235788018 | C | T | 0.296 | 0.161 | 0.032 | 3.06E-07 | 282266 | AGAP1-IT1 |
| rs12472150 | 2 | 235788047 | G | A | 0.296 | 0.161 | 0.032 | 3.06E-07 | 282295 | AGAP1-IT1 |
| rs56329604 | 2 | 235788220 | T | C | 0.296 | 0.161 | 0.032 | 3.06E-07 | 282468 | AGAP1-IT1 |
| rs72983003 | 2 | 235788415 | T | G | 0.296 | 0.161 | 0.032 | 3.06E-07 | 282663 | AGAP1-IT1 |
| rs72983005 | 2 | 235788471 | C | T | 0.296 | 0.161 | 0.032 | 3.06E-07 | 282719 | AGAP1-IT1 |
| rs72983007 | 2 | 235788621 | G | T | 0.296 | 0.161 | 0.032 | 3.06E-07 | 282869 | AGAP1-IT1 |
| rs72983009 | 2 | 235788684 | G | A | 0.296 | 0.161 | 0.032 | 3.06E-07 | 282932 | AGAP1-IT1 |
| rs72983012 | 2 | 235788851 | T | C | 0.296 | 0.161 | 0.032 | 3.06E-07 | 283099 | AGAP1-IT1 |
| rs12469161 | 2 | 235789283 | G | C | 0.296 | 0.161 | 0.032 | 3.06E-07 | 283531 | AGAP1-IT1 |
| rs12469185 | 2 | 235789305 | T | C | 0.296 | 0.161 | 0.032 | 3.06E-07 | 283553 | AGAP1-IT1 |
| rs12474276 | 2 | 235789568 | A | T | 0.296 | 0.161 | 0.032 | 3.06E-07 | 283816 | AGAP1-IT1 |
| rs56062711 | 2 | 235789772 | T | G | 0.296 | 0.161 | 0.032 | 3.06E-07 | 284020 | AGAP1-IT1 |
| rs55936237 | 2 | 235789838 | T | C | 0.296 | 0.161 | 0.032 | 3.06E-07 | 284086 | AGAP1-IT1 |
| rs72983019 | 2 | 235790229 | C | T | 0.296 | 0.161 | 0.032 | 3.06E-07 | 284477 | AGAP1-IT1 |
| rs12475524 | 2 | 235790749 | G | T | 0.296 | 0.161 | 0.032 | 3.06E-07 | 284997 | AGAP1-IT1 |
| rs12474606 | 2 | 235790763 | G | A | 0.296 | 0.161 | 0.032 | 3.06E-07 | 285011 | AGAP1-IT1 |
| rs12470496 | 2 | 235790785 | T | C | 0.296 | 0.161 | 0.032 | 3.06E-07 | 285033 | AGAP1-IT1 |
| rs12474630 | 2 | 235790852 | G | A | 0.306 | 0.161 | 0.032 | 4.11E-07 | 285100 | AGAP1-IT1 |
| rs55881123 | 2 | 235790935 | C | T | 0.296 | 0.161 | 0.032 | 3.06E-07 | 285183 | AGAP1-IT1 |
| rs72983030 | 2 | 235791262 | G | A | 0.296 | 0.161 | 0.032 | 3.06E-07 | 285510 | AGAP1-IT1 |
| rs72983032 | 2 | 235791271 | C | T | 0.296 | 0.161 | 0.032 | 3.06E-07 | 285519 | AGAP1-IT1 |
| rs6845244 | 4 | 14785233 | C | A | 0.806 | -0.177 | 0.038 | 2.74E-06 | 102937 | LINC00504 |
| rs4698291 | 4 | 14788174 | G | C | 0.806 | -0.177 | 0.038 | 2.74E-06 | 99996 | LINC00504 |
| rs13151601 | 4 | 27958744 | T | A | 0.724 | 0.170 | 0.038 | 6.30E-06 | 741264 | LINC02261 |
| rs7665009 | 4 | 27960206 | T | C | 0.724 | 0.170 | 0.038 | 6.30E-06 | 742726 | LINC02261 |
| rs7438137 | 4 | 75831883 | T | C | 0.439 | 0.145 | 0.032 | 5.02E-06 | 70689 | PPEF2 |
| rs62326190 | 4 | 155956020 | C | T | 0.194 | 0.176 | 0.038 | 3.24E-06 | 197027 | GUCY1B1 |
| rs4306914 | 4 | 156020957 | G | C | 0.194 | 0.176 | 0.038 | 3.24E-06 | 261964 | GUCY1B1 |
| rs371103786 | 4 | 181401805 | T | C | 0.286 | 0.183 | 0.040 | 4.71E-06 | -242655 | LINC00290 |
| rs4113603 | 4 | 184286837 | C | T | 0.347 | -0.170 | 0.037 | 4.85E-06 | -65606 | ENPP6 |
| rs1217971 | 4 | 184288728 | T | C | 0.622 | 0.180 | 0.038 | 2.63E-06 | 65250 | LINC02363 |
| rs1217972 | 4 | 184288886 | G | T | 0.622 | 0.180 | 0.038 | 2.63E-06 | 65092 | LINC02363 |
| rs6453850 | 6 | 75937483 | A | G | 0.306 | 0.170 | 0.036 | 1.84E-06 | 135196 | IMPG1 |
| rs9341546 | 6 | 75937859 | T | C | 0.653 | -0.157 | 0.033 | 2.49E-06 | 134820 | IMPG1 |
| rs9350607 | 6 | 75937890 | C | T | 0.653 | -0.157 | 0.033 | 2.49E-06 | 134789 | IMPG1 |
| rs9360962 | 6 | 75938923 | G | T | 0.663 | -0.165 | 0.035 | 1.85E-06 | 133756 | IMPG1 |
| rs9343338 | 6 | 75939209 | T | A | 0.663 | -0.165 | 0.035 | 1.85E-06 | 133470 | IMPG1 |
| rs13207399 | 6 | 75939355 | G | C | 0.306 | 0.170 | 0.036 | 1.84E-06 | 133324 | IMPG1 |
| rs2313752 | 6 | 75940342 | C | T | 0.663 | -0.165 | 0.035 | 1.85E-06 | 132337 | IMPG1 |
| rs2313753 | 6 | 75942546 | G | C | 0.663 | -0.165 | 0.035 | 1.85E-06 | 130133 | IMPG1 |
| rs2313755 | 6 | 75943204 | T | C | 0.673 | -0.173 | 0.033 | 2.03E-07 | 129475 | IMPG1 |
| rs1610336 | 6 | 75943378 | A | G | 0.306 | 0.170 | 0.036 | 1.84E-06 | 129301 | IMPG1 |
| rs76014128 | 6 | 78481469 | C | T | 0.163 | 0.209 | 0.044 | 2.46E-06 | -386004 | IRAK1BP1 |
| rs76364903 | 6 | 78482432 | A | G | 0.184 | 0.199 | 0.044 | 5.69E-06 | -385041 | IRAK1BP1 |
| rs11968517 | 6 | 78486543 | C | G | 0.163 | 0.209 | 0.044 | 2.46E-06 | -380930 | IRAK1BP1 |
| rs997648 | 6 | 78488462 | A | G | 0.163 | 0.209 | 0.044 | 2.46E-06 | -379011 | IRAK1BP1 |
| rs78808189 | 6 | 78489069 | A | T | 0.163 | 0.209 | 0.044 | 2.46E-06 | -378404 | IRAK1BP1 |
| rs77972979 | 6 | 78489165 | T | C | 0.163 | 0.209 | 0.044 | 2.46E-06 | -378308 | IRAK1BP1 |
| rs79700878 | 6 | 78489720 | A | G | 0.163 | 0.209 | 0.044 | 2.46E-06 | -377753 | IRAK1BP1 |
| rs1407103 | 6 | 78491257 | A | C | 0.163 | 0.209 | 0.044 | 2.46E-06 | -376216 | IRAK1BP1 |
| rs7773443 | 6 | 78494501 | A | G | 0.163 | 0.209 | 0.044 | 2.46E-06 | -372972 | IRAK1BP1 |
| rs76167616 | 6 | 78494757 | T | C | 0.163 | 0.209 | 0.044 | 2.46E-06 | -372716 | IRAK1BP1 |
| rs76312719 | 6 | 78495068 | C | T | 0.163 | 0.209 | 0.044 | 2.46E-06 | -372405 | IRAK1BP1 |
| rs77974320 | 6 | 78495334 | C | T | 0.163 | 0.209 | 0.044 | 2.46E-06 | -372139 | IRAK1BP1 |

| | | | | | | | | | | |
|---|---|---|---|---|---|---|---|---|---|---|
| rs148823079 | 6 | 78495409 | C | T | 0.163 | 0.209 | 0.044 | 2.46E-06 | -372064 | IRAK1BP1 |
| rs114251753 | 6 | 78495612 | A | G | 0.163 | 0.209 | 0.044 | 2.46E-06 | -371861 | IRAK1BP1 |
| rs80099930 | 6 | 78495713 | C | T | 0.163 | 0.209 | 0.044 | 2.46E-06 | -371760 | IRAK1BP1 |
| rs115355509 | 6 | 78495767 | G | A | 0.163 | 0.209 | 0.044 | 2.46E-06 | -371706 | IRAK1BP1 |
| rs200194409 | 6 | 78495969 | A | C | 0.163 | 0.209 | 0.044 | 2.46E-06 | -371504 | IRAK1BP1 |
| rs11961883 | 6 | 78496288 | A | G | 0.163 | 0.209 | 0.044 | 2.46E-06 | -371185 | IRAK1BP1 |
| rs7751932 | 6 | 78497915 | A | C | 0.163 | 0.209 | 0.044 | 2.46E-06 | -369558 | IRAK1BP1 |
| rs79949695 | 6 | 78499776 | C | G | 0.163 | 0.209 | 0.044 | 2.46E-06 | -367697 | IRAK1BP1 |
| rs7744184 | 6 | 78501326 | A | G | 0.163 | 0.209 | 0.044 | 2.46E-06 | -366147 | IRAK1BP1 |
| rs79313931 | 6 | 78504256 | A | G | 0.163 | 0.209 | 0.044 | 2.46E-06 | -363217 | IRAK1BP1 |
| rs6900179 | 6 | 78507704 | C | G | 0.163 | 0.209 | 0.044 | 2.46E-06 | -359769 | IRAK1BP1 |
| rs6922709 | 6 | 78507743 | C | T | 0.163 | 0.209 | 0.044 | 2.46E-06 | -359730 | IRAK1BP1 |
| rs1884966 | 6 | 78508721 | A | G | 0.163 | 0.209 | 0.044 | 2.46E-06 | -358752 | IRAK1BP1 |
| rs60520348 | 6 | 78510501 | G | T | 0.163 | 0.209 | 0.044 | 2.46E-06 | -356972 | IRAK1BP1 |
| rs11967020 | 6 | 78514356 | T | C | 0.163 | 0.209 | 0.044 | 2.46E-06 | -353117 | IRAK1BP1 |
| rs1358817 | 6 | 78518080 | A | C | 0.163 | 0.209 | 0.044 | 2.46E-06 | -349393 | IRAK1BP1 |
| rs723437 | 6 | 78521437 | G | A | 0.163 | 0.209 | 0.044 | 2.46E-06 | -346036 | IRAK1BP1 |
| rs9359328 | 6 | 78524372 | A | G | 0.163 | 0.209 | 0.044 | 2.46E-06 | -343101 | IRAK1BP1 |
| rs9343777 | 6 | 78525400 | G | T | 0.163 | 0.209 | 0.044 | 2.46E-06 | -342073 | IRAK1BP1 |
| rs1321597 | 6 | 78527929 | T | C | 0.163 | 0.209 | 0.044 | 2.46E-06 | -339544 | IRAK1BP1 |
| rs76641149 | 6 | 78541449 | A | C | 0.143 | 0.237 | 0.043 | 4.60E-08 | -326024 | IRAK1BP1 |
| rs79747327 | 6 | 78556991 | G | A | 0.133 | 0.221 | 0.046 | 1.88E-06 | -310482 | IRAK1BP1 |
| rs76889818 | 6 | 78566330 | A | G | 0.133 | 0.221 | 0.046 | 1.88E-06 | -301143 | IRAK1BP1 |
| rs77413528 | 6 | 78581094 | G | A | 0.133 | 0.221 | 0.046 | 1.88E-06 | -286379 | IRAK1BP1 |
| rs9359652 | 6 | 85344324 | A | G | 0.418 | 0.151 | 0.034 | 8.55E-06 | 45863 | LINC02535 |
| rs12201128 | 6 | 85353689 | G | T | 0.418 | 0.151 | 0.034 | 8.55E-06 | 36498 | LINC02535 |
| rs11771342 | 7 | 46204239 | T | G | 0.163 | 0.216 | 0.044 | 7.61E-07 | -282364 | IGFBP3 |
| rs9691259 | 7 | 46206982 | G | A | 0.163 | 0.216 | 0.044 | 7.61E-07 | -285107 | IGFBP3 |
| rs2633359 | 7 | 138122716 | C | T | 0.888 | -0.250 | 0.055 | 4.64E-06 | -1043 | MIR4468 |
| rs2440798 | 7 | 138127010 | T | C | 0.888 | -0.250 | 0.055 | 4.64E-06 | 3251 | MIR4468 |
| rs4368897 | 7 | 146986807 | T | C | 0.153 | 0.205 | 0.046 | 6.51E-06 | 110803 | CNTNAP2-AS1 |
| rs12682543 | 8 | 29222041 | A | G | 0.643 | -0.153 | 0.034 | 8.36E-06 | 41084 | KIF13B |
| rs13263888 | 8 | 29223321 | C | G | 0.643 | -0.153 | 0.034 | 8.36E-06 | 39804 | KIF13B |
| rs4585791 | 9 | 614530 | A | T | 0.235 | -0.197 | 0.042 | 2.68E-06 | 144238 | KANK1 |
| rs11812737 | 10 | 62669559 | A | G | 0.173 | 0.200 | 0.040 | 5.15E-07 | -135299 | ADO |
| rs4746518 | 10 | 62669835 | G | A | 0.214 | 0.171 | 0.038 | 6.35E-06 | -135023 | ADO |
| rs729738 | 10 | 62670505 | A | C | 0.214 | 0.171 | 0.038 | 6.35E-06 | -134353 | ADO |
| rs729739 | 10 | 62670542 | A | G | 0.214 | 0.171 | 0.038 | 6.35E-06 | -134316 | ADO |
| rs7394165 | 10 | 62677258 | T | C | 0.173 | 0.200 | 0.040 | 5.15E-07 | -127600 | ADO |
| rs7393302 | 10 | 62677294 | G | T | 0.214 | 0.171 | 0.038 | 6.35E-06 | -127564 | ADO |
| rs4746524 | 10 | 62678037 | T | C | 0.173 | 0.200 | 0.040 | 5.15E-07 | -126821 | ADO |
| rs224121 | 10 | 62687592 | C | A | 0.827 | -0.216 | 0.043 | 3.96E-07 | -117266 | ADO |
| rs2960666 | 10 | 115819198 | G | A | 0.673 | -0.156 | 0.035 | 8.81E-06 | 454270 | GFRA1 |
| rs372086626 | 11 | 39730859 | C | T | 0.173 | 0.208 | 0.044 | 1.73E-06 | 1082000 | LINC01493 |
| rs11063601 | 12 | 562434 | G | T | 0.112 | 0.248 | 0.055 | 6.25E-06 | 24053 | LOC105369595 |
| rs3217810 | 12 | 4279105 | T | C | 0.153 | 0.205 | 0.046 | 6.45E-06 | -2920 | CCND2-AS1 |
| rs2075267 | 12 | 14962167 | G | T | 0.612 | -0.150 | 0.032 | 2.84E-06 | -438 | ARHGDIB |
| rs11056258 | 12 | 14964465 | T | G | 0.592 | -0.156 | 0.033 | 1.70E-06 | -2736 | ARHGDIB |
| rs7311989 | 12 | 14964603 | A | C | 0.592 | -0.156 | 0.033 | 1.70E-06 | -2874 | ARHGDIB |
| rs7300467 | 12 | 14964663 | G | T | 0.602 | -0.148 | 0.033 | 6.58E-06 | -2934 | ARHGDIB |
| rs10846097 | 12 | 14970137 | T | C | 0.643 | -0.161 | 0.036 | 7.28E-06 | -2886 | PDE6H |
| rs7978415 | 12 | 14970534 | A | G | 0.592 | -0.156 | 0.033 | 1.70E-06 | -2489 | PDE6H |
| rs3983696 | 12 | 14970938 | T | C | 0.592 | -0.156 | 0.033 | 1.70E-06 | -2085 | PDE6H |
| rs7313555 | 12 | 14988129 | C | T | 0.663 | -0.158 | 0.035 | 8.00E-06 | -13705 | LINC01489 |
| rs11058862 | 12 | 122592803 | T | C | 0.398 | 0.141 | 0.032 | 7.77E-06 | 65556 | KNTC1 |
| rs11611668 | 12 | 122609181 | T | C | 0.398 | 0.143 | 0.031 | 4.87E-06 | 81934 | KNTC1 |
| rs61947369 | 13 | 25920538 | A | G | 0.163 | 0.186 | 0.042 | 9.46E-06 | 130494 | SHISA2 |

| | | | | | | | | | | |
|---|---|---|---|---|---|---|---|---|---|---|
| rs9548368 | 13 | 38482144 | T | C | 0.367 | 0.151 | 0.033 | 4.62E-06 | 63156 | LINC00437 |
| rs9532237 | 13 | 38482716 | A | G | 0.367 | 0.151 | 0.033 | 4.62E-06 | 62584 | LINC00437 |
| rs6571996 | 14 | 40299024 | A | C | 0.745 | -0.184 | 0.036 | 3.61E-07 | -655875 | LINC02315 |
| rs7160617 | 14 | 40326534 | G | A | 0.806 | -0.217 | 0.047 | 3.19E-06 | -628365 | LINC02315 |
| rs6494413 | 15 | 63564443 | A | G | 0.755 | -0.161 | 0.036 | 6.55E-06 | -32911 | FBXL22 |
| rs72625754 | 15 | 63564562 | C | T | 0.173 | 0.207 | 0.039 | 1.20E-07 | -32792 | FBXL22 |
| rs6494414 | 15 | 63566507 | G | A | 0.755 | -0.161 | 0.036 | 6.55E-06 | -30847 | FBXL22 |
| rs6494415 | 15 | 63570130 | T | G | 0.755 | -0.161 | 0.036 | 6.55E-06 | -27224 | FBXL22 |
| rs8028681 | 15 | 63572006 | T | C | 0.755 | -0.161 | 0.036 | 6.55E-06 | -25348 | FBXL22 |
| rs8029310 | 15 | 63572221 | T | G | 0.755 | -0.161 | 0.036 | 6.55E-06 | -25133 | FBXL22 |
| rs7183892 | 15 | 63586733 | T | C | 0.755 | -0.161 | 0.036 | 6.55E-06 | -10621 | FBXL22 |
| rs7168622 | 15 | 63668675 | A | C | 0.122 | 0.230 | 0.047 | 9.81E-07 | -67085 | USP3-AS1 |
| rs7182375 | 15 | 63813431 | G | A | 0.776 | -0.163 | 0.036 | 4.85E-06 | 20512 | HERC1 |
| rs7170689 | 15 | 63831046 | T | C | 0.776 | -0.159 | 0.036 | 9.59E-06 | 2897 | HERC1 |
| rs8042418 | 15 | 63832686 | A | G | 0.776 | -0.159 | 0.036 | 9.59E-06 | 1257 | HERC1 |
| rs7170433 | 15 | 63837452 | C | T | 0.776 | -0.159 | 0.036 | 9.59E-06 | -3509 | HERC1 |
| rs214239 | 16 | 303625 | A | G | 0.214 | 0.182 | 0.040 | 5.50E-06 | 20472 | PDIA2 |
| rs1558346 | 16 | 6514333 | T | C | 0.143 | 0.208 | 0.046 | 7.22E-06 | 881865 | MIR8065 |
| rs72804582 | 16 | 86720970 | G | A | 0.112 | 0.246 | 0.055 | 8.44E-06 | 985 | LINC02189 |
| rs11659939 | 18 | 33845019 | A | G | 0.214 | 0.179 | 0.040 | 9.37E-06 | 266403 | ASXL3 |
| rs1051651 | 19 | 7116272 | C | G | 0.245 | 0.166 | 0.035 | 2.23E-06 | 46827 | ZNF557 |
| rs11670340 | 19 | 7436914 | A | G | 0.408 | 0.138 | 0.031 | 7.47E-06 | 54079 | ARHGEF18 |
| rs6139617 | 20 | 5063227 | G | A | 0.469 | -0.143 | 0.031 | 2.93E-06 | 49877 | TMEM230 |

**Table S6.** Annotation of variants associated with GO_ATP_METABOLIC level with p-val<$10^{-5}$.

| rsID | CHR | POS | A1 | A2 | AF1 | BETA | SE | P | distancetoFeature | symbol |
|---|---|---|---|---|---|---|---|---|---|---|
| rs4649064 | 1 | 25094616 | G | A | 0.456 | 0.232 | 0.045 | 2.07E-07 | 71112 | RUNX1 |
| rs6657249 | 1 | 25095126 | C | A | 0.395 | 0.250 | 0.044 | 1.65E-08 | 71622 | RUNX1 |
| rs11588172 | 1 | 25095499 | C | T | 0.395 | 0.250 | 0.044 | 1.65E-08 | 71995 | RUNX1 |
| rs56365408 | 1 | 25096844 | G | A | 0.395 | 0.250 | 0.044 | 1.65E-08 | 73340 | RUNX1 |
| rs56348607 | 1 | 25097067 | G | A | 0.412 | 0.258 | 0.045 | 1.23E-08 | 73563 | RUNX1 |
| rs58693652 | 1 | 25097185 | A | G | 0.395 | 0.250 | 0.044 | 1.65E-08 | 73681 | RUNX1 |
| rs2536363 | 1 | 83382963 | C | T | 0.728 | -0.254 | 0.056 | 6.46E-06 | -396754 | LINC01361 |
| rs2536361 | 1 | 83383171 | T | G | 0.728 | -0.254 | 0.056 | 6.46E-06 | -396962 | LINC01361 |
| N.A | 1 | 121043250 | C | G | 0.728 | -0.201 | 0.041 | 1.25E-06 | -44096 | FCGR1B |
| rs12471349 | 2 | 231630391 | A | C | 0.456 | -0.203 | 0.043 | 2.12E-06 | 37489 | TEX44 |
| rs11686124 | 2 | 231631051 | A | G | 0.447 | -0.200 | 0.042 | 2.28E-06 | 38149 | TEX44 |
| rs12619551 | 2 | 231632587 | T | C | 0.447 | -0.200 | 0.042 | 2.28E-06 | 39685 | TEX44 |
| rs12621259 | 2 | 231633982 | T | C | 0.447 | -0.200 | 0.042 | 2.28E-06 | 41080 | TEX44 |
| rs10204481 | 2 | 231635421 | T | C | 0.456 | -0.203 | 0.043 | 2.12E-06 | 42519 | TEX44 |
| rs10204963 | 2 | 231635773 | A | G | 0.456 | -0.203 | 0.043 | 2.12E-06 | 42871 | TEX44 |
| rs11685588 | 2 | 231635997 | A | C | 0.456 | -0.203 | 0.043 | 2.12E-06 | 43095 | TEX44 |
| rs931390 | 3 | 4736178 | T | C | 0.746 | -0.269 | 0.051 | 1.26E-07 | 15413 | EGOT |
| rs4690284 | 4 | 73616 | C | T | 0.105 | 0.315 | 0.071 | 9.38E-06 | 20330 | ZNF595 |
| rs11735004 | 4 | 8519493 | A | G | 0.105 | 0.359 | 0.077 | 3.52E-06 | -58833 | CPZ |
| rs28520722 | 4 | 168372678 | C | G | 0.158 | 0.261 | 0.055 | 2.56E-06 | -53870 | DDX60 |
| rs10474554 | 5 | 78503335 | T | C | 0.816 | -0.256 | 0.054 | 2.01E-06 | 142751 | SCAMP1 |
| rs10474555 | 5 | 78503370 | T | C | 0.816 | -0.256 | 0.054 | 2.01E-06 | 142786 | SCAMP1 |
| rs12522528 | 5 | 172824478 | C | A | 0.272 | 0.247 | 0.049 | 5.68E-07 | -9798 | ERGIC1 |
| rs6891132 | 5 | 172824828 | G | A | 0.711 | -0.224 | 0.049 | 4.38E-06 | -9448 | ERGIC1 |
| rs4867689 | 5 | 172825091 | C | T | 0.711 | -0.224 | 0.049 | 4.38E-06 | -9185 | ERGIC1 |
| rs4867690 | 5 | 172825370 | G | A | 0.711 | -0.224 | 0.049 | 4.38E-06 | -8906 | ERGIC1 |
| rs4868216 | 5 | 172825449 | A | G | 0.711 | -0.224 | 0.049 | 4.38E-06 | -8827 | ERGIC1 |
| rs4867691 | 5 | 172825540 | G | C | 0.711 | -0.224 | 0.049 | 4.38E-06 | -8736 | ERGIC1 |
| rs7717831 | 5 | 172825774 | C | T | 0.711 | -0.224 | 0.049 | 4.38E-06 | -8502 | ERGIC1 |
| rs7717986 | 5 | 172825824 | A | T | 0.711 | -0.224 | 0.049 | 4.38E-06 | -8452 | ERGIC1 |
| rs7736149 | 5 | 172825871 | A | G | 0.711 | -0.224 | 0.049 | 4.38E-06 | -8405 | ERGIC1 |
| rs7736270 | 5 | 172825917 | A | G | 0.711 | -0.224 | 0.049 | 4.38E-06 | -8359 | ERGIC1 |
| rs792994 | 5 | 172826179 | T | C | 0.272 | 0.247 | 0.049 | 5.68E-07 | -8097 | ERGIC1 |
| rs11748400 | 5 | 172826590 | G | T | 0.263 | 0.255 | 0.049 | 1.53E-07 | -7686 | ERGIC1 |
| rs4868217 | 5 | 172826780 | C | T | 0.711 | -0.224 | 0.049 | 4.38E-06 | -7496 | ERGIC1 |
| rs12519115 | 5 | 172827357 | T | C | 0.254 | 0.259 | 0.048 | 7.76E-08 | -6919 | ERGIC1 |
| rs7707991 | 5 | 172827470 | G | A | 0.711 | -0.224 | 0.049 | 4.38E-06 | -6806 | ERGIC1 |
| rs6556047 | 5 | 172828846 | A | T | 0.702 | -0.211 | 0.047 | 8.90E-06 | -5430 | ERGIC1 |
| rs1564259 | 5 | 172829021 | C | T | 0.702 | -0.211 | 0.047 | 8.90E-06 | -5255 | ERGIC1 |
| rs1006721 | 5 | 172829068 | A | G | 0.702 | -0.211 | 0.047 | 8.90E-06 | -5208 | ERGIC1 |
| rs10057822 | 5 | 172829748 | G | A | 0.702 | -0.211 | 0.047 | 8.90E-06 | -4528 | ERGIC1 |
| rs2339652 | 5 | 172830696 | G | A | 0.702 | -0.211 | 0.047 | 8.90E-06 | -3580 | ERGIC1 |
| rs4868218 | 5 | 172830968 | A | C | 0.702 | -0.211 | 0.047 | 8.90E-06 | -3308 | ERGIC1 |
| rs35566639 | 5 | 172834459 | A | G | 0.254 | 0.250 | 0.049 | 3.70E-07 | 183 | ERGIC1 |
| rs35768025 | 5 | 172834486 | A | C | 0.254 | 0.250 | 0.049 | 3.70E-07 | 210 | ERGIC1 |
| rs78078909 | 5 | 172834535 | A | C | 0.254 | 0.250 | 0.049 | 3.70E-07 | 259 | ERGIC1 |
| rs793010 | 5 | 172835670 | T | C | 0.263 | 0.241 | 0.050 | 1.31E-06 | 1394 | ERGIC1 |
| rs6895746 | 5 | 172838337 | G | A | 0.254 | 0.263 | 0.048 | 3.69E-08 | 4061 | ERGIC1 |
| rs11134762 | 5 | 172838397 | A | G | 0.237 | 0.267 | 0.047 | 1.87E-08 | 4121 | ERGIC1 |

| | | | | | | | | | | |
|---|---|---|---|---|---|---|---|---|---|---|
| rs11949146 | 5 | 172840207 | A | G | 0.219 | 0.282 | 0.050 | 1.54E-08 | 5931 | ERGIC1 |
| rs6877332 | 5 | 172841627 | A | G | 0.246 | 0.267 | 0.051 | 1.61E-07 | 7351 | ERGIC1 |
| rs7722970 | 5 | 172842868 | T | G | 0.219 | 0.258 | 0.052 | 8.18E-07 | 8592 | ERGIC1 |
| rs10945708 | 6 | 161017963 | T | C | 0.298 | 0.227 | 0.049 | 3.32E-06 | 26235 | MAP3K4 |
| rs12195182 | 6 | 161108526 | C | G | 0.465 | 0.195 | 0.042 | 4.49E-06 | 53457 | AGPAT4-IT1 |
| rs2314157 | 6 | 161190126 | C | G | 0.439 | 0.194 | 0.042 | 3.99E-06 | -28143 | AGPAT4-IT1 |
| rs3735487 | 7 | 45065547 | G | A | 0.114 | 0.274 | 0.060 | 4.18E-06 | 23368 | NACAD |
| rs2140953 | 7 | 106848675 | C | A | 0.105 | 0.354 | 0.078 | 5.32E-06 | -16604 | PIK3CG |
| rs7791674 | 7 | 106849681 | T | C | 0.105 | 0.354 | 0.078 | 5.32E-06 | -15598 | PIK3CG |
| rs35031873 | 7 | 142758607 | C | A | 0.079 | 0.385 | 0.074 | 2.17E-07 | -373471 | PIP |
| rs1567924 | 8 | 19201161 | G | C | 0.237 | 0.244 | 0.053 | 4.21E-06 | 58309 | LOC100128993 |
| rs62524263 | 8 | 102126785 | G | C | 0.649 | 0.214 | 0.047 | 5.51E-06 | 1352 | MIR5680 |
| rs6982041 | 8 | 102127205 | C | A | 0.649 | 0.214 | 0.047 | 5.51E-06 | 1772 | MIR5680 |
| rs7819114 | 8 | 102128424 | T | C | 0.649 | 0.214 | 0.047 | 5.51E-06 | 2991 | MIR5680 |
| rs4734053 | 8 | 102129163 | A | G | 0.649 | 0.214 | 0.047 | 5.51E-06 | 3730 | MIR5680 |
| rs4734616 | 8 | 102129293 | T | C | 0.658 | 0.215 | 0.046 | 3.56E-06 | 3860 | MIR5680 |
| rs7826028 | 8 | 102131357 | C | T | 0.649 | 0.214 | 0.047 | 5.51E-06 | 5924 | MIR5680 |
| rs7000087 | 8 | 102131841 | G | T | 0.640 | 0.218 | 0.047 | 3.59E-06 | 6408 | MIR5680 |
| rs9942801 | 8 | 102132950 | G | A | 0.649 | 0.214 | 0.047 | 5.51E-06 | 7517 | MIR5680 |
| rs4734617 | 8 | 102138783 | C | A | 0.649 | 0.214 | 0.047 | 5.51E-06 | 13350 | MIR5680 |
| rs13248776 | 8 | 102145359 | T | C | 0.649 | 0.214 | 0.047 | 5.51E-06 | 19926 | MIR5680 |
| rs13250882 | 8 | 102145366 | C | T | 0.649 | 0.214 | 0.047 | 5.51E-06 | 19933 | MIR5680 |
| rs6988832 | 8 | 102146011 | G | A | 0.649 | 0.214 | 0.047 | 5.51E-06 | 20578 | MIR5680 |
| rs2105616 | 8 | 102146140 | C | A | 0.649 | 0.214 | 0.047 | 5.51E-06 | 20707 | MIR5680 |
| rs372431986 | 8 | 102146141 | C | A | 0.649 | 0.214 | 0.047 | 5.51E-06 | 20708 | MIR5680 |
| rs1892965 | 8 | 102146620 | A | C | 0.649 | 0.214 | 0.047 | 5.51E-06 | 21187 | MIR5680 |
| rs13268773 | 8 | 102147224 | A | G | 0.649 | 0.214 | 0.047 | 5.51E-06 | 21791 | MIR5680 |
| rs6468817 | 8 | 102148609 | A | T | 0.649 | 0.214 | 0.047 | 5.51E-06 | 23176 | MIR5680 |
| rs12681037 | 8 | 102149097 | A | T | 0.649 | 0.214 | 0.047 | 5.51E-06 | 23664 | MIR5680 |
| rs6468818 | 8 | 102149937 | G | C | 0.649 | 0.214 | 0.047 | 5.51E-06 | 24504 | MIR5680 |
| rs4291239 | 8 | 102150410 | A | G | 0.649 | 0.214 | 0.047 | 5.51E-06 | 24977 | MIR5680 |
| rs10093024 | 8 | 102151656 | C | T | 0.649 | 0.214 | 0.047 | 5.51E-06 | 26223 | MIR5680 |
| rs6992948 | 8 | 102152160 | A | C | 0.596 | 0.220 | 0.045 | 1.27E-06 | 26727 | MIR5680 |
| rs1111911 | 8 | 102152888 | T | A | 0.649 | 0.214 | 0.047 | 5.51E-06 | 27455 | MIR5680 |
| rs6980613 | 8 | 102153223 | G | T | 0.649 | 0.214 | 0.047 | 5.51E-06 | 27790 | MIR5680 |
| rs1573311 | 8 | 102154787 | T | C | 0.649 | 0.214 | 0.047 | 5.51E-06 | 29354 | MIR5680 |
| rs2387095 | 8 | 102155415 | G | T | 0.649 | 0.214 | 0.047 | 5.51E-06 | 29982 | MIR5680 |
| rs1547370 | 8 | 102155949 | T | C | 0.649 | 0.214 | 0.047 | 5.51E-06 | 30516 | MIR5680 |
| rs2186682 | 8 | 102157973 | G | A | 0.649 | 0.214 | 0.047 | 5.51E-06 | 32540 | MIR5680 |
| rs1573309 | 8 | 102160658 | G | A | 0.605 | 0.234 | 0.049 | 1.48E-06 | 35225 | MIR5680 |
| rs1148514 | 8 | 102168709 | C | G | 0.605 | 0.234 | 0.049 | 1.48E-06 | 43276 | MIR5680 |
| rs1148515 | 8 | 102168828 | G | C | 0.605 | 0.234 | 0.049 | 1.48E-06 | 43395 | MIR5680 |
| rs1148516 | 8 | 102168999 | G | A | 0.605 | 0.234 | 0.049 | 1.48E-06 | 43566 | MIR5680 |
| rs78848775 | 8 | 133734878 | T | A | 0.105 | 0.319 | 0.071 | 6.46E-06 | 42475 | LOC101927798 |
| rs77131790 | 8 | 133735142 | G | A | 0.105 | 0.319 | 0.071 | 6.46E-06 | 42211 | LOC101927798 |
| rs75149129 | 8 | 133735347 | G | T | 0.105 | 0.319 | 0.071 | 6.46E-06 | 42006 | LOC101927798 |
| rs76106243 | 8 | 133737227 | C | T | 0.105 | 0.319 | 0.071 | 6.46E-06 | 40126 | LOC101927798 |
| rs17820827 | 9 | 13863723 | G | C | 0.167 | 0.286 | 0.062 | 4.62E-06 | -64249 | LINC00583 |
| rs7858214 | 9 | 13865261 | T | C | 0.105 | 0.327 | 0.070 | 3.09E-06 | -62711 | LINC00583 |
| rs11259468 | 10 | 15081387 | C | G | 0.114 | 0.308 | 0.069 | 9.07E-06 | 7390 | ACBD7 |

| | | | | | | | | | | |
|---|---|---|---|---|---|---|---|---|---|---|
| rs10901547 | 10 | 126174101 | T | C | 0.281 | 0.236 | 0.048 | 7.53E-07 | -200974 | FANK1-AS1 |
| rs72826371 | 10 | 126177069 | T | C | 0.237 | 0.256 | 0.049 | 1.45E-07 | -203942 | FANK1-AS1 |
| rs12356272 | 10 | 126177590 | G | A | 0.272 | 0.242 | 0.047 | 2.49E-07 | -204463 | FANK1-AS1 |
| rs12357824 | 10 | 126180060 | C | T | 0.272 | 0.242 | 0.047 | 2.49E-07 | -206933 | FANK1-AS1 |
| rs12766940 | 10 | 126180562 | C | T | 0.272 | 0.242 | 0.047 | 2.49E-07 | -207435 | FANK1-AS1 |
| rs11244871 | 10 | 126183597 | A | G | 0.263 | 0.249 | 0.046 | 6.57E-08 | 204859 | ADAM12 |
| rs1459709 | 10 | 126184259 | A | G | 0.263 | 0.249 | 0.046 | 6.57E-08 | 204197 | ADAM12 |
| rs17683203 | 10 | 126186844 | A | G | 0.263 | 0.249 | 0.046 | 6.57E-08 | 201612 | ADAM12 |
| rs17745507 | 10 | 126187170 | T | A | 0.228 | 0.265 | 0.048 | 2.93E-08 | 201286 | ADAM12 |
| rs10829685 | 10 | 130058387 | T | G | 0.228 | 0.259 | 0.052 | 6.16E-07 | 52431 | C10orf143 |
| rs7082237 | 10 | 130091533 | T | C | 0.140 | 0.272 | 0.061 | 8.76E-06 | 19285 | C10orf143 |
| rs1945331 | 11 | 21242296 | T | C | 0.123 | 0.304 | 0.068 | 7.66E-06 | 572744 | NELL1 |
| rs4755920 | 11 | 44947205 | A | G | 0.360 | -0.214 | 0.048 | 6.66E-06 | 4102 | TP53I11 |
| rs141434045 | 12 | 115147278 | G | A | 0.123 | 0.352 | 0.072 | 1.08E-06 | -463113 | TBX5-TBX3 |
| rs10082870 | 12 | 115148523 | C | G | 0.123 | 0.352 | 0.072 | 1.08E-06 | -464358 | TBX5-TBX3 |
| rs2270458 | 12 | 115151267 | T | C | 0.114 | 0.364 | 0.074 | 8.36E-07 | -467102 | TBX5-TBX3 |
| rs12310526 | 12 | 115151871 | A | G | 0.114 | 0.364 | 0.074 | 8.36E-07 | -467706 | TBX5-TBX3 |
| rs112088312 | 12 | 115154065 | T | C | 0.114 | 0.364 | 0.074 | 8.36E-07 | -469900 | TBX5-TBX3 |
| rs11067442 | 12 | 115154219 | T | C | 0.114 | 0.364 | 0.074 | 8.36E-07 | -470054 | TBX5-TBX3 |
| rs74846266 | 12 | 115154979 | T | C | 0.114 | 0.364 | 0.074 | 8.36E-07 | -470814 | TBX5-TBX3 |
| rs7970502 | 12 | 115155723 | T | C | 0.114 | 0.364 | 0.074 | 8.36E-07 | -471558 | TBX5-TBX3 |
| rs7145901 | 14 | 52220916 | T | C | 0.228 | 0.251 | 0.057 | 9.57E-06 | -46798 | PTGDR |
| rs71474496 | 15 | 42625943 | G | A | 0.123 | 0.318 | 0.067 | 1.90E-06 | 50283 | STARD9 |
| rs11638835 | 15 | 42626143 | G | A | 0.868 | -0.301 | 0.066 | 5.41E-06 | 50483 | STARD9 |
| rs12911585 | 15 | 42642347 | C | T | 0.132 | 0.306 | 0.066 | 3.31E-06 | 66687 | STARD9 |
| rs61122145 | 15 | 42699689 | A | G | 0.149 | 0.306 | 0.063 | 1.12E-06 | 37438 | CDAN1 |
| rs12902830 | 15 | 42705184 | T | C | 0.140 | 0.320 | 0.063 | 3.93E-07 | 31943 | CDAN1 |
| rs35392023 | 15 | 42709462 | A | G | 0.149 | 0.311 | 0.062 | 6.02E-07 | 27665 | CDAN1 |
| rs1058846 | 15 | 42719826 | T | C | 0.149 | 0.311 | 0.062 | 6.02E-07 | 17301 | CDAN1 |
| rs17774047 | 15 | 42721028 | T | C | 0.149 | 0.311 | 0.062 | 6.02E-07 | 16099 | CDAN1 |
| rs35088691 | 15 | 42772565 | C | T | 0.140 | 0.328 | 0.062 | 1.39E-07 | -35438 | CDAN1 |
| rs12913618 | 15 | 42779131 | T | C | 0.140 | 0.328 | 0.062 | 1.39E-07 | -42004 | CDAN1 |
| rs12915116 | 15 | 42782482 | T | C | 0.140 | 0.328 | 0.062 | 1.39E-07 | -45355 | CDAN1 |
| rs35298309 | 15 | 42784803 | C | T | 0.149 | 0.337 | 0.060 | 1.86E-08 | -47676 | CDAN1 |
| rs12908349 | 15 | 42794029 | T | C | 0.123 | 0.315 | 0.067 | 2.59E-06 | -56902 | CDAN1 |
| rs66932813 | 15 | 42796424 | G | T | 0.123 | 0.315 | 0.067 | 2.59E-06 | -59297 | CDAN1 |
| rs36092914 | 15 | 42808425 | A | C | 0.123 | 0.315 | 0.067 | 2.59E-06 | -71298 | CDAN1 |
| rs964776412 | 15 | 42817281 | A | T | 0.123 | 0.315 | 0.067 | 2.59E-06 | -80154 | CDAN1 |
| rs72713784 | 15 | 42823738 | G | A | 0.123 | 0.315 | 0.067 | 2.59E-06 | -86611 | CDAN1 |
| rs34745458 | 15 | 42845003 | C | T | 0.123 | 0.315 | 0.067 | 2.59E-06 | 75807 | TTBK2 |
| rs72713795 | 15 | 42849430 | A | T | 0.123 | 0.315 | 0.067 | 2.59E-06 | 71380 | TTBK2 |
| rs35202249 | 15 | 42855844 | T | C | 0.123 | 0.315 | 0.067 | 2.59E-06 | 64966 | TTBK2 |
| rs2683240 | 15 | 81120333 | T | C | 0.675 | -0.209 | 0.046 | 5.84E-06 | -62268 | IL16 |
| rs2460852 | 15 | 81123862 | T | G | 0.675 | -0.209 | 0.046 | 5.84E-06 | -58739 | IL16 |
| rs2683249 | 15 | 81129249 | G | A | 0.675 | -0.209 | 0.046 | 5.84E-06 | -53352 | IL16 |
| rs2683255 | 15 | 81134198 | C | G | 0.675 | -0.209 | 0.046 | 5.84E-06 | -48403 | IL16 |
| rs2683256 | 15 | 81135816 | A | G | 0.675 | -0.209 | 0.046 | 5.84E-06 | -46785 | IL16 |
| rs4404021 | 15 | 96060577 | T | C | 0.228 | 0.227 | 0.048 | 2.70E-06 | -265362 | NR2F2 |
| rs7166802 | 15 | 96062205 | T | G | 0.237 | 0.219 | 0.049 | 7.62E-06 | -263734 | NR2F2 |
| rs7166314 | 15 | 96062218 | G | A | 0.237 | 0.219 | 0.049 | 7.62E-06 | -263721 | NR2F2 |

| | | | | | | | | | | |
|---|---|---|---|---|---|---|---|---|---|---|
| rs79651776 | 16 | 11650396 | G | A | 0.140 | 0.273 | 0.061 | 7.50E-06 | -14014 | LITAF |
| rs8056041 | 16 | 83121479 | A | G | 0.263 | 0.240 | 0.053 | 6.68E-06 | -131180 | LOC101928417 |
| rs8082569 | 17 | 62828928 | G | A | 0.491 | 0.198 | 0.045 | 9.81E-06 | -20583 | MARCHF10 |
| rs12948372 | 17 | 62831387 | A | G | 0.412 | 0.211 | 0.047 | 6.96E-06 | -23042 | MARCHF10 |
| rs35594211 | 17 | 62831620 | G | A | 0.412 | 0.211 | 0.047 | 6.96E-06 | -23275 | MARCHF10 |
| rs11868614 | 17 | 62831835 | A | G | 0.412 | 0.211 | 0.047 | 6.96E-06 | -23490 | MARCHF10 |
| rs35354702 | 17 | 62832130 | G | A | 0.412 | 0.211 | 0.047 | 6.96E-06 | -23785 | MARCHF10 |
| rs35596881 | 17 | 62834618 | G | A | 0.412 | 0.211 | 0.047 | 6.96E-06 | -26273 | MARCHF10 |
| rs8182298 | 17 | 62837616 | C | T | 0.439 | 0.208 | 0.043 | 1.08E-06 | -29271 | MARCHF10 |
| rs8182316 | 17 | 62837746 | T | C | 0.456 | 0.207 | 0.042 | 1.13E-06 | -29401 | MARCHF10 |
| rs8076190 | 17 | 62837908 | G | A | 0.456 | 0.207 | 0.042 | 1.13E-06 | -29563 | MARCHF10 |
| rs8077260 | 17 | 62838375 | C | G | 0.456 | 0.207 | 0.042 | 1.13E-06 | -30030 | MARCHF10 |
| rs8182324 | 17 | 62838599 | G | A | 0.456 | 0.207 | 0.042 | 1.13E-06 | -30254 | MARCHF10 |
| rs6504130 | 17 | 62838901 | T | C | 0.421 | 0.215 | 0.043 | 4.92E-07 | -30556 | MARCHF10 |
| rs8064710 | 17 | 62839334 | C | T | 0.439 | 0.208 | 0.043 | 1.08E-06 | -30989 | MARCHF10 |
| rs8064402 | 17 | 62839705 | A | G | 0.439 | 0.208 | 0.043 | 1.08E-06 | -31360 | MARCHF10 |
| rs8067960 | 17 | 62839798 | T | C | 0.439 | 0.208 | 0.043 | 1.08E-06 | -31453 | MARCHF10 |
| rs7226368 | 17 | 62840771 | C | G | 0.421 | 0.215 | 0.043 | 4.92E-07 | -32426 | MARCHF10 |
| rs12943672 | 17 | 62841241 | A | T | 0.439 | 0.208 | 0.043 | 1.08E-06 | -32896 | MARCHF10 |
| rs1476811 | 17 | 62841947 | T | C | 0.421 | 0.215 | 0.043 | 4.92E-07 | -33602 | MARCHF10 |
| rs1476812 | 17 | 62842115 | T | C | 0.421 | 0.215 | 0.043 | 4.92E-07 | -33770 | MARCHF10 |
| rs1987647 | 17 | 62842528 | C | T | 0.377 | 0.216 | 0.046 | 2.90E-06 | -34183 | MARCHF10 |
| rs12943559 | 17 | 62843136 | T | C | 0.421 | 0.215 | 0.043 | 4.92E-07 | -34791 | MARCHF10 |
| rs9903891 | 17 | 62843788 | A | T | 0.447 | 0.213 | 0.043 | 8.16E-07 | -35443 | MARCHF10 |
| rs12936567 | 17 | 62844192 | A | G | 0.447 | 0.213 | 0.043 | 8.16E-07 | -35847 | MARCHF10 |
| rs11871628 | 17 | 62844788 | C | T | 0.447 | 0.213 | 0.043 | 8.16E-07 | -36443 | MARCHF10 |
| rs35395730 | 17 | 62845085 | C | A | 0.447 | 0.213 | 0.043 | 8.16E-07 | -36740 | MARCHF10 |
| rs55895685 | 17 | 62846360 | C | G | 0.439 | 0.214 | 0.044 | 1.50E-06 | -38015 | MARCHF10 |
| rs55854611 | 17 | 62846443 | G | T | 0.439 | 0.214 | 0.044 | 1.50E-06 | -38098 | MARCHF10 |
| rs55909719 | 17 | 62846445 | A | G | 0.404 | 0.210 | 0.044 | 1.85E-06 | -38100 | MARCHF10 |
| rs8072675 | 17 | 62846458 | G | A | 0.439 | 0.214 | 0.044 | 1.50E-06 | -38113 | MARCHF10 |
| rs34350332 | 17 | 62847043 | C | T | 0.439 | 0.214 | 0.044 | 1.50E-06 | -38698 | MARCHF10 |
| rs34826284 | 17 | 62847771 | T | C | 0.404 | 0.230 | 0.044 | 2.15E-07 | -39426 | MARCHF10 |
| rs35290531 | 17 | 62849438 | A | C | 0.439 | 0.214 | 0.044 | 1.50E-06 | -41093 | MARCHF10 |
| rs35572014 | 17 | 62849602 | C | T | 0.447 | 0.213 | 0.045 | 2.75E-06 | -41257 | MARCHF10 |
| rs12051594 | 17 | 62850242 | A | G | 0.456 | 0.212 | 0.044 | 1.49E-06 | -41897 | MARCHF10 |
| rs9915826 | 17 | 62850877 | T | C | 0.421 | 0.215 | 0.045 | 1.71E-06 | -42532 | MARCHF10 |
| rs9303464 | 17 | 62850993 | C | T | 0.447 | 0.213 | 0.045 | 2.75E-06 | -42648 | MARCHF10 |
| rs9890310 | 17 | 62851050 | G | T | 0.447 | 0.213 | 0.045 | 2.75E-06 | -42705 | MARCHF10 |
| rs9890570 | 17 | 62851197 | C | T | 0.421 | 0.215 | 0.045 | 1.71E-06 | -42852 | MARCHF10 |
| rs8076646 | 17 | 62852550 | T | C | 0.386 | 0.209 | 0.045 | 3.44E-06 | -44205 | MARCHF10 |
| rs8081770 | 17 | 62852855 | C | T | 0.465 | 0.217 | 0.047 | 3.98E-06 | -44510 | MARCHF10 |
| rs61638683 | 17 | 82849592 | T | G | 0.535 | -0.210 | 0.045 | 3.38E-06 | -9013 | ZNF750 |
| rs59300995 | 17 | 82849594 | C | G | 0.553 | -0.206 | 0.044 | 2.47E-06 | -9015 | ZNF750 |
| rs7216324 | 17 | 82851589 | C | A | 0.535 | -0.210 | 0.045 | 3.38E-06 | -11010 | ZNF750 |
| rs4986118 | 17 | 82851689 | C | G | 0.447 | 0.206 | 0.044 | 2.47E-06 | -11110 | ZNF750 |
| rs7215939 | 17 | 82851732 | G | C | 0.535 | -0.210 | 0.045 | 3.38E-06 | -11153 | ZNF750 |
| rs7214952 | 17 | 82851853 | T | G | 0.535 | -0.210 | 0.045 | 3.38E-06 | -11274 | ZNF750 |
| rs11077948 | 17 | 82852057 | T | C | 0.553 | -0.206 | 0.044 | 2.47E-06 | -11478 | ZNF750 |
| rs9913679 | 17 | 82855022 | G | A | 0.535 | -0.210 | 0.045 | 3.38E-06 | -14443 | ZNF750 |

| | | | | | | | | | | |
|---|---|---|---|---|---|---|---|---|---|---|
| rs12600507 | 17 | 82856293 | A | G | 0.553 | -0.206 | 0.044 | 2.47E-06 | -15714 | ZNF750 |
| rs9303013 | 17 | 82857421 | C | T | 0.544 | -0.212 | 0.046 | 4.73E-06 | -16842 | ZNF750 |
| rs7220732 | 17 | 82857901 | C | A | 0.544 | -0.212 | 0.046 | 4.73E-06 | -17322 | ZNF750 |
| rs7209378 | 17 | 82859192 | A | G | 0.561 | -0.208 | 0.045 | 3.36E-06 | -18613 | ZNF750 |
| rs9890196 | 17 | 82860297 | C | T | 0.553 | -0.213 | 0.045 | 2.58E-06 | -19718 | ZNF750 |
| rs6502007 | 17 | 82861401 | G | A | 0.544 | -0.196 | 0.043 | 6.08E-06 | -20822 | ZNF750 |
| rs8082007 | 17 | 82862130 | A | G | 0.526 | -0.210 | 0.046 | 5.71E-06 | -21551 | ZNF750 |
| rs8065192 | 17 | 82862158 | G | A | 0.553 | -0.206 | 0.044 | 2.47E-06 | -21579 | ZNF750 |
| rs9303014 | 17 | 82866493 | T | G | 0.526 | -0.210 | 0.046 | 5.71E-06 | -25914 | ZNF750 |
| rs4986120 | 17 | 82866778 | G | T | 0.535 | -0.204 | 0.046 | 7.70E-06 | -26199 | ZNF750 |
| rs7224427 | 17 | 82867331 | A | G | 0.526 | -0.210 | 0.046 | 5.71E-06 | -26752 | ZNF750 |
| rs56353584 | 17 | 82869089 | G | T | 0.553 | -0.207 | 0.046 | 6.21E-06 | -28510 | ZNF750 |
| rs56322462 | 17 | 82869216 | T | C | 0.544 | -0.212 | 0.046 | 4.73E-06 | -28637 | ZNF750 |
| rs66893622 | 17 | 82869369 | T | A | 0.535 | -0.212 | 0.047 | 8.08E-06 | -28790 | ZNF750 |
| rs9892064 | 17 | 82870027 | T | C | 0.544 | -0.212 | 0.046 | 4.73E-06 | -29448 | ZNF750 |
| rs3744161 | 17 | 82870181 | A | G | 0.535 | -0.212 | 0.047 | 8.08E-06 | -29602 | ZNF750 |
| rs6502008 | 17 | 82872389 | A | G | 0.544 | -0.212 | 0.046 | 4.73E-06 | -31810 | ZNF750 |
| rs6502009 | 17 | 82872499 | T | C | 0.544 | -0.212 | 0.046 | 4.73E-06 | -31920 | ZNF750 |
| rs11655504 | 17 | 82873448 | C | T | 0.535 | -0.212 | 0.047 | 8.08E-06 | -32869 | ZNF750 |
| rs898093 | 17 | 82874591 | T | C | 0.535 | -0.212 | 0.047 | 8.08E-06 | -34012 | ZNF750 |
| rs4986123 | 17 | 82876370 | T | C | 0.544 | -0.205 | 0.045 | 4.52E-06 | -35791 | ZNF750 |
| rs8075190 | 17 | 82878558 | T | C | 0.544 | -0.201 | 0.045 | 8.17E-06 | -37979 | ZNF750 |
| rs9747095 | 17 | 82882712 | G | A | 0.474 | 0.204 | 0.044 | 4.30E-06 | -42133 | ZNF750 |
| rs7221830 | 17 | 82890938 | G | T | 0.474 | 0.204 | 0.044 | 4.30E-06 | -50359 | ZNF750 |
| rs59103510 | 17 | 82891072 | T | G | 0.518 | -0.204 | 0.045 | 7.08E-06 | -50493 | ZNF750 |
| rs12948880 | 17 | 82893224 | C | T | 0.474 | 0.210 | 0.046 | 5.71E-06 | -52645 | ZNF750 |
| rs4986130 | 17 | 82895802 | C | T | 0.482 | 0.204 | 0.045 | 7.08E-06 | -55223 | ZNF750 |
| rs4986131 | 17 | 82895920 | G | A | 0.482 | 0.204 | 0.045 | 7.08E-06 | -55341 | ZNF750 |
| rs12951357 | 17 | 82896533 | A | G | 0.474 | 0.210 | 0.046 | 5.71E-06 | -55954 | ZNF750 |
| rs1809252 | 17 | 82897473 | A | G | 0.518 | -0.204 | 0.045 | 7.08E-06 | -56894 | ZNF750 |
| rs939262 | 17 | 82897656 | G | C | 0.518 | -0.204 | 0.045 | 7.08E-06 | -57077 | ZNF750 |
| rs9891862 | 17 | 82900793 | A | G | 0.518 | -0.204 | 0.045 | 7.08E-06 | -60214 | ZNF750 |
| rs4986134 | 17 | 82901278 | A | G | 0.518 | -0.204 | 0.045 | 7.08E-06 | -60699 | ZNF750 |
| rs2292971 | 17 | 82905981 | C | T | 0.482 | 0.204 | 0.045 | 7.08E-06 | -65402 | ZNF750 |
| rs8074371 | 17 | 82908853 | C | G | 0.482 | 0.204 | 0.045 | 7.08E-06 | -68274 | ZNF750 |
| rs12947897 | 17 | 82913585 | G | A | 0.474 | 0.204 | 0.044 | 4.16E-06 | -73006 | ZNF750 |
| rs8064687 | 17 | 82915798 | C | G | 0.482 | 0.204 | 0.045 | 7.08E-06 | -75219 | ZNF750 |
| rs8069773 | 17 | 82916667 | T | G | 0.474 | 0.204 | 0.044 | 4.16E-06 | -76088 | ZNF750 |
| rs73007122 | 19 | 10004217 | T | C | 0.105 | 0.338 | 0.069 | 1.08E-06 | 6255 | COL5A3 |
| rs35143140 | 19 | 10005321 | A | G | 0.289 | 0.218 | 0.047 | 2.95E-06 | 5151 | COL5A3 |
| rs11085530 | 19 | 10006027 | G | A | 0.289 | 0.218 | 0.047 | 2.95E-06 | 4445 | COL5A3 |
| rs35678764 | 19 | 10006948 | A | C | 0.289 | 0.218 | 0.047 | 2.95E-06 | 3524 | COL5A3 |
| rs9797822 | 19 | 10007058 | A | C | 0.289 | 0.218 | 0.047 | 2.95E-06 | 3414 | COL5A3 |
| rs9797907 | 19 | 10007075 | G | T | 0.289 | 0.218 | 0.047 | 2.95E-06 | 3397 | COL5A3 |
| rs11672609 | 19 | 58495557 | T | C | 0.140 | 0.258 | 0.057 | 6.77E-06 | 16509 | SLC27A5 |
| rs11672614 | 19 | 58495617 | T | C | 0.140 | 0.258 | 0.057 | 6.77E-06 | 16449 | SLC27A5 |
| rs11672730 | 19 | 58495836 | A | G | 0.140 | 0.258 | 0.057 | 6.77E-06 | 16230 | SLC27A5 |
| rs55652736 | 19 | 58496199 | T | G | 0.140 | 0.258 | 0.057 | 6.77E-06 | 15867 | SLC27A5 |
| rs73066226 | 19 | 58496846 | C | T | 0.149 | 0.256 | 0.057 | 6.21E-06 | 15220 | SLC27A5 |
| rs11671092 | 19 | 58497742 | C | T | 0.149 | 0.256 | 0.057 | 6.21E-06 | 14324 | SLC27A5 |

| | | | | | | | | | | |
|---|---|---|---|---|---|---|---|---|---|---|
| rs55928441 | 19 | 58498307 | C | T | 0.140 | 0.258 | 0.057 | 6.77E-06 | 13759 | SLC27A5 |
| rs73066228 | 19 | 58501829 | G | A | 0.140 | 0.258 | 0.057 | 7.22E-06 | 10237 | SLC27A5 |
| rs4810126 | 20 | 58244817 | T | C | 0.167 | 0.298 | 0.061 | 1.23E-06 | -16163 | ANKRD60 |
| rs76912023 | 20 | 58257517 | T | C | 0.158 | 0.300 | 0.062 | 1.39E-06 | -28863 | ANKRD60 |
| rs4811976 | 20 | 58259149 | A | G | 0.158 | 0.300 | 0.062 | 1.39E-06 | -30495 | ANKRD60 |
| rs563489 | 20 | 58355773 | G | A | 0.649 | -0.237 | 0.051 | 3.08E-06 | -33350 | VAPB |
| rs71315381 | 22 | 17510895 | G | C | 0.140 | 0.274 | 0.061 | 6.67E-06 | -52545 | SLC25A18 |
| rs2142664 | 22 | 46559201 | G | A | 0.237 | 0.227 | 0.051 | 8.47E-06 | -16812 | GRAMD4 |

**Table S7**. Annotation of variants associated with *GATA4* activity level with p-val<$10^{-5}$.
